# Systematic identification of chromatin organizers as tuners of intratumoral heterogeneity

**DOI:** 10.64898/2026.04.18.719392

**Authors:** Brian J Woo, Sushil Sobti, Jung Min Suh, Hassan Yousefi, Kristle Garcia, Shaopu Zhou, Ashir Borah, Hani Goodarzi

## Abstract

Intratumoral transcriptional heterogeneity (ITH) enables tumor cells to explore diverse phenotypic states, fueling therapeutic resistance and metastatic progression. Yet systematic approaches for identifying non-genetic regulators of this phenotype have been lacking. Here, we develop a multi-scale screening framework that integrates bulk patient transcriptomics, functional *in vivo* genetic screening, and single-cell transcriptomic and epigenomic profiling to nominate chromatin organizers that modulate ITH in breast cancer. From an initial set of 41 candidates nominated *in silico*, we identify RNF8 and MIS18A as factors whose perturbation produces coordinated shifts in transcriptomic variability and chromatin accessibility dispersion at the single-cell level, without altering mean expression or genomic copy number. MIS18A acts through a centromere-independent, DNA-methylation-associated pathway. Modulating RNF8 or MIS18A expression tunes chemotherapeutic sensitivity and metastatic potential *in vivo*: knockdown sensitizes cells and reduces metastatic burden, whereas overexpression confers resistance and promotes progression. Higher expression of both genes predicts shorter patient survival. These findings identify RNF8 and MIS18A as chromatin-level stochastic tuners of transcriptional variability in cancer and provide a generalizable framework for discovering regulators of non-genetic tumor heterogeneity.

## 1. Introduction

Intratumoral heterogeneity—cell-to-cell variation within a single tumor—is a central driver of tumor progression, therapeutic resistance, and recurrence ^1–7^. Although targeted molecular and immunological therapies can initially reduce tumor burden, the emergence of resistant subpopulations from a heterogeneous tumor almost invariably leads to relapse and mortality ^8–11^. Increased transcriptional plasticity, in particular, enables tumor subpopulations to access varied transcriptomic states and thereby survive in unfamiliar microenvironments, including those imposed by chemotherapeutic stress ^12–15^. Once a subpopulation reaches a transcriptomic state that enhances fitness, it is selected for and propagated, driving the growth of resistant tumor populations.

Intratumoral transcriptional heterogeneity (ITH) can arise through both genetic and non-genetic mechanisms ^16–26^. Genetic sources, including localized mutations and focal amplifications, have been extensively characterized ^14,15,27,28^. Non-genetic mechanisms, however, remain less well understood, despite mounting evidence that they contribute substantially to ITH and to clinically relevant phenotypes such as drug resistance ^29–34^. We previously showed that cancer subpopulations exhibiting enhanced transcriptome variability display increased metastatic fitness in breast cancer ^12^, motivating a deeper investigation of the molecular regulators that govern this variability.

A candidate mechanism has recently emerged from studies of stochastic tuning; a process whereby inherent transcriptional noise enables cells to explore gene expression states that confer adaptive advantages. First demonstrated in yeast ^13^, stochastic tuning was recently extended to mammalian cells adapting to chemotherapeutic challenge ^35^. Critically, both studies implicated chromatin-modification machinery as a modulator of this process, suggesting that chromatin organizers (COs) may act as regulators of transcriptional heterogeneity. In the context of cancer, stochastic tuning could allow tumors to explore transcriptomic states that confer resistance to previously unencountered selective pressures such as hypoxia or drug exposure ^11,12^. We propose that the capacity to expand the total range of searchable transcriptomic states, which is governed in part by chromatin organizers, represents a general mechanism underlying tumor progression.

To identify COs that regulate ITH, we required a strategy that could nominate candidates from existing large-scale datasets. Sufficiently powered single-cell datasets linking CO expression to cell-to-cell transcriptomic variability are not yet available at the scale needed for systematic discovery. We therefore reasoned that, based on the central limit theorem, factors that increase cell-to-cell transcriptomic variability within tumors should also increase the variance of bulk expression measurements across tumors. While this signal will be attenuated by within-tumor averaging, it may nonetheless be detectable given the thousands of bulk tumor profiles that are available. Using this heuristic, we nominated chromatin organizers whose expression was associated with increased transcriptomic variability among breast cancer samples. Intertumoral variability does not necessarily imply intratumoral heterogeneity (i.e., variability within individual tumors), however, it can serve as a tractable proxy for initial candidate nomination.

By integrating this *in silico* nomination with *in vivo* functional genetic screening for tumor progression and single-cell transcriptomic, morphological, and epigenomic profiling, we converged on two chromatin organizers previously uncharacterized in the context of ITH: RNF8 and MIS18A ^36–41^. We showed that RNF8 tunes TWIST1-directed EMT, while MIS18A modulates heterogeneity through DNMT3B-dependent changes in genomic methylation state. Perturbation of either factor altered chemotherapeutic sensitivity and tumor fitness *in vivo*, and expression of both correlated with survival in breast cancer patient cohorts.

## 2. Results

### 2.1. *In silico* nomination of chromatin organizers associated with transcriptomic variability

We sought to identify chromatin organizers (COs) that modulate intratumoral transcriptional heterogeneity (ITH). As outlined above, existing single-cell breast cancer cohorts lack the sample size and perturbation coverage needed for systematic, genome-wide discovery of ITH regulators. We therefore used bulk TCGA breast cancer RNA-seq data as an *in silico* nomination strategy, reasoning that COs driving cell-to-cell transcriptomic variability within tumors should also increase transcriptomic variability across patients (see Methods). For each CO (drawn from Gene Ontology annotations of human chromatin organizers ^42^), we stratified patients by expression level and tested whether genome-wide transcript coefficient of variation (CV) differed significantly between high- and low-expressing groups, benchmarked against an empirical null distribution derived from random patient groupings (**Figure 1A**, **S1A**, Supplementary Tables 1, 2).

**Figure 1.**
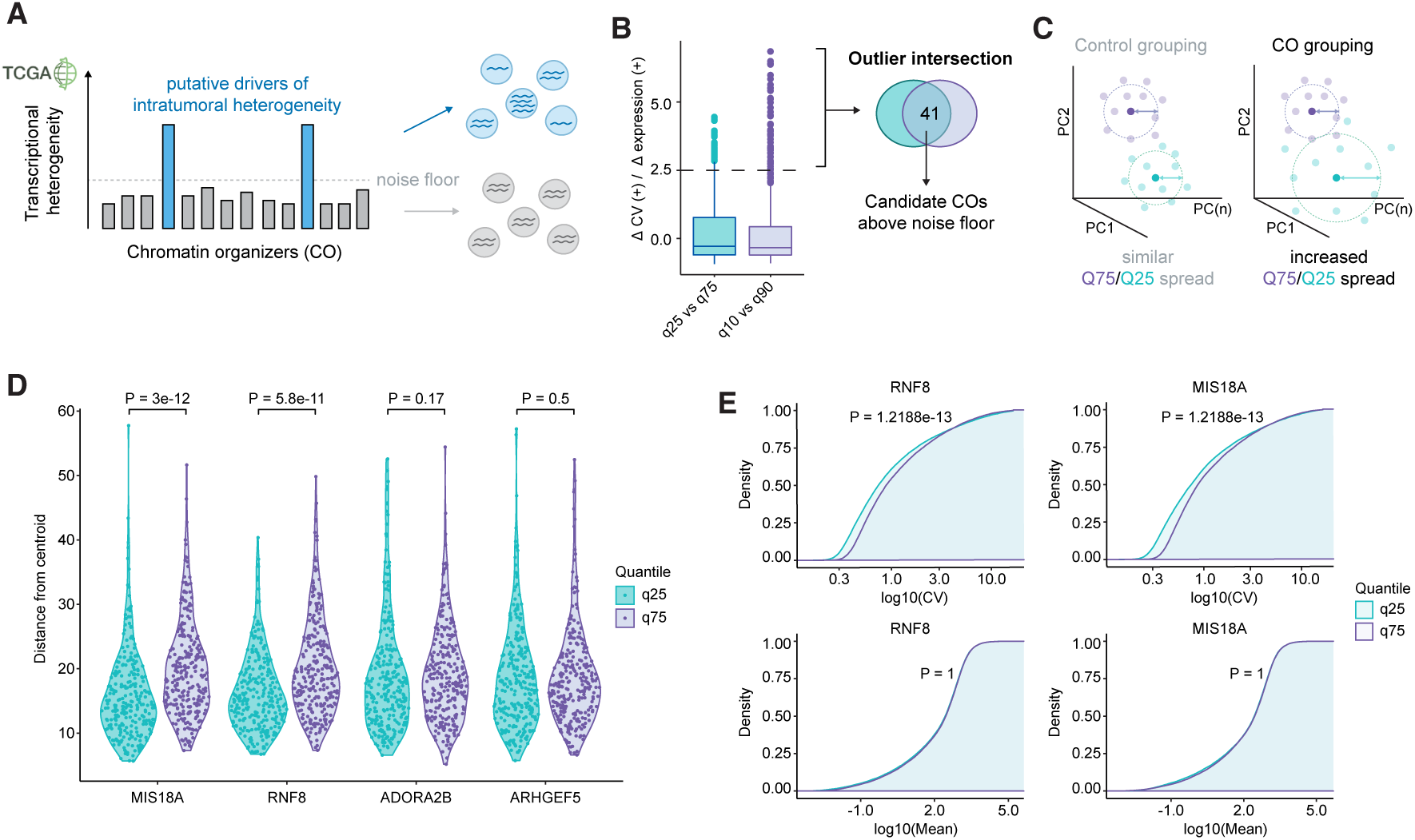
*In silico* nomination of chromatin organizers associated with increased transcriptomic variability. **(A)** Schematic of the computational pipeline. TCGA breast cancer RNA-seq patients were stratified by expression of each chromatin organizer (CO), and genome-wide transcript coefficient of variation (CV) was compared between high- and low-expressing groups. An empirical noise floor was established using random patient groupings (*N* = 50). COs with signal above this threshold were nominated as candidate regulators of transcriptomic heterogeneity. **(B)** Nominated COs from quartile and quintile stratifications. Each point represents one CO; axes show the change in CV (ΔCV) and the empirical *z*-score for each stratification scheme. 41 COs satisfied the *z* > 2.5 cutoff in both analyses. **(C)** Schematic illustrating the PCA-based spread metric. For each patient group, the distance from each patient’s transcriptomic profile to the group centroid was computed across the top 50 principal components. Control groupings (left) show similar spread between quartiles; CO-based groupings (right) show increased spread in the high-expression quartile. **(D)** Distance from quartile transcriptomic centroid for two representative COs (RNF8 and MIS18A, left) and two negative control genes (right). *P*-values by two-sided Wilcoxon signed-rank test. **(E)** CV density distributions (top) and mean expression density distributions (bottom) for representative CO quartile groupings. The rightward shift in CV was not accompanied by a corresponding shift in mean expression, indicating that increased variability is not driven by changes in expression magnitude. *P*-values by two-sided Kolmogorov–Smirnov test, adjusted with the Benjamini–Hochberg procedure.

This analysis nominated 41 COs whose elevated expression was associated with increased intertumoral transcriptomic variability across both quartile and quintile stratification schemes (**Figure 1B**). To visualize this effect, we projected patient transcriptomes into principal component space and found that patients in the top CO expression quartile showed greater transcriptomic spread than those in the bottom quartile, relative to uncorrelated control groupings (**Figure 1C–D**, **S1B**). We next asked whether the increased transcriptomic CV could be explained by accompanying changes in genomic copy number, given the well-documented role of copy number amplification (CNA) and aneuploidy in driving transcriptomic heterogeneity ^1,6–8,10^. Genes with significant transcriptomic CV changes were uncorrelated with those showing significant CNA changes, suggesting that the observed variability operates through non-genetic mechanisms (**Figure S1C**). Consistent with this, increased transcriptomic heterogeneity was not accompanied by a global shift in mean transcript abundance (**Figure 1E**), and tumor purity scores remained high across patient groupings, ruling out confounding by shifts in cellular composition (**Figure S1D**).

### 2.2. *In vivo* CRISPRi screening identifies COs that drive tumor progression

Because drivers of transcriptomic heterogeneity are also expected to promote tumor progression ^12^, we next sought to functionally filter our 41 nominated COs for those with a direct impact on tumor fitness. We performed an *in vivo* CRISPRi ^43^ screen in the MDA-MB-231 breast cancer xenograft model, targeting all 41 COs with a pooled sgRNA library (∼5 guides per CO; see Methods). We assayed both orthotopic mammary fat pad injection (primary tumor progression) and tail vein injection (metastatic lung colonization), comparing sgRNA representation in tumors to cells grown *in vitro* for a matched number of doublings (**Figure 2A**). Of the 41 COs, 16 showed *in vivo*-specific depletion upon CRISPRi-mediated silencing across both screening modalities, indicating that their expression is required for tumor fitness (**Figure 2B**, Supplementary Tables 3, 4).

**Figure 2.**
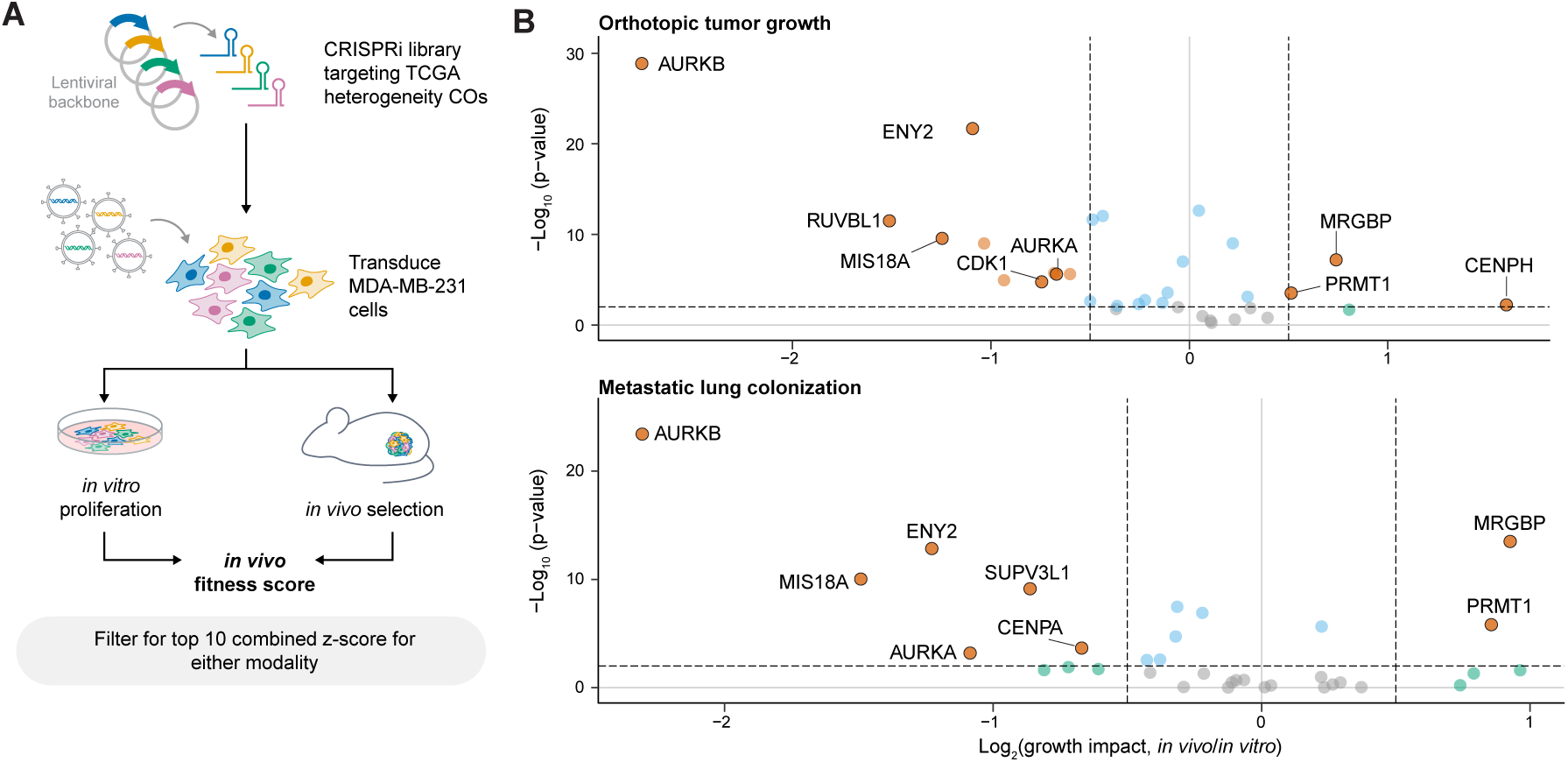
*In vivo* CRISPRi screen identifies chromatin organizers required for tumor progression. **(A)** Schematic of the pooled CRISPRi screen. MDA-MB-231 cells transduced with a CO-targeting sgRNA library were injected via orthotopic (mammary fat pad) or tail-vein routes into female NSG mice. sgRNA representation was compared between *in vivo* tumors and size-matched *in vitro* controls. **(B)** Volcano plots of screen results for orthotopic injection (left) and tail-vein injection (right). The *x*-axis shows fitness scores (sgRNA representation in tumor normalized to *in vitro* endpoint); positive values indicate increased representation *in vivo*. The *y*-axis shows − log_10_ (*P*).

### 2.3. RNF8 and MIS18A modulate transcriptomic heterogeneity at the single-cell level

To determine whether these 16 COs are required and sufficient to influence transcriptomic heterogeneity at the single-cell level, we performed CRISPRi and CRISPRa Perturb-seq ^43,44^ targeting all 16 COs in MDA-MB-231 cells (**Figure 3A**; see Methods). We calculated genome-wide CV for each perturbation and compared it to size-matched random control groupings. Fewer than half of the 16 COs modulated transcriptomic variability in this assay (**Figure S2A–C**); notably, AURKB showed discordant behavior relative to its bulk TCGA signal, likely due to broad secondary transcriptomic effects of perturbing an essential mitotic regulator. Intersecting the CRISPRi and CRISPRa results, RNF8 and MIS18A emerged as the two strongest modulators of single-cell transcriptomic variability (**Figure 3B**).

**Figure 3.**
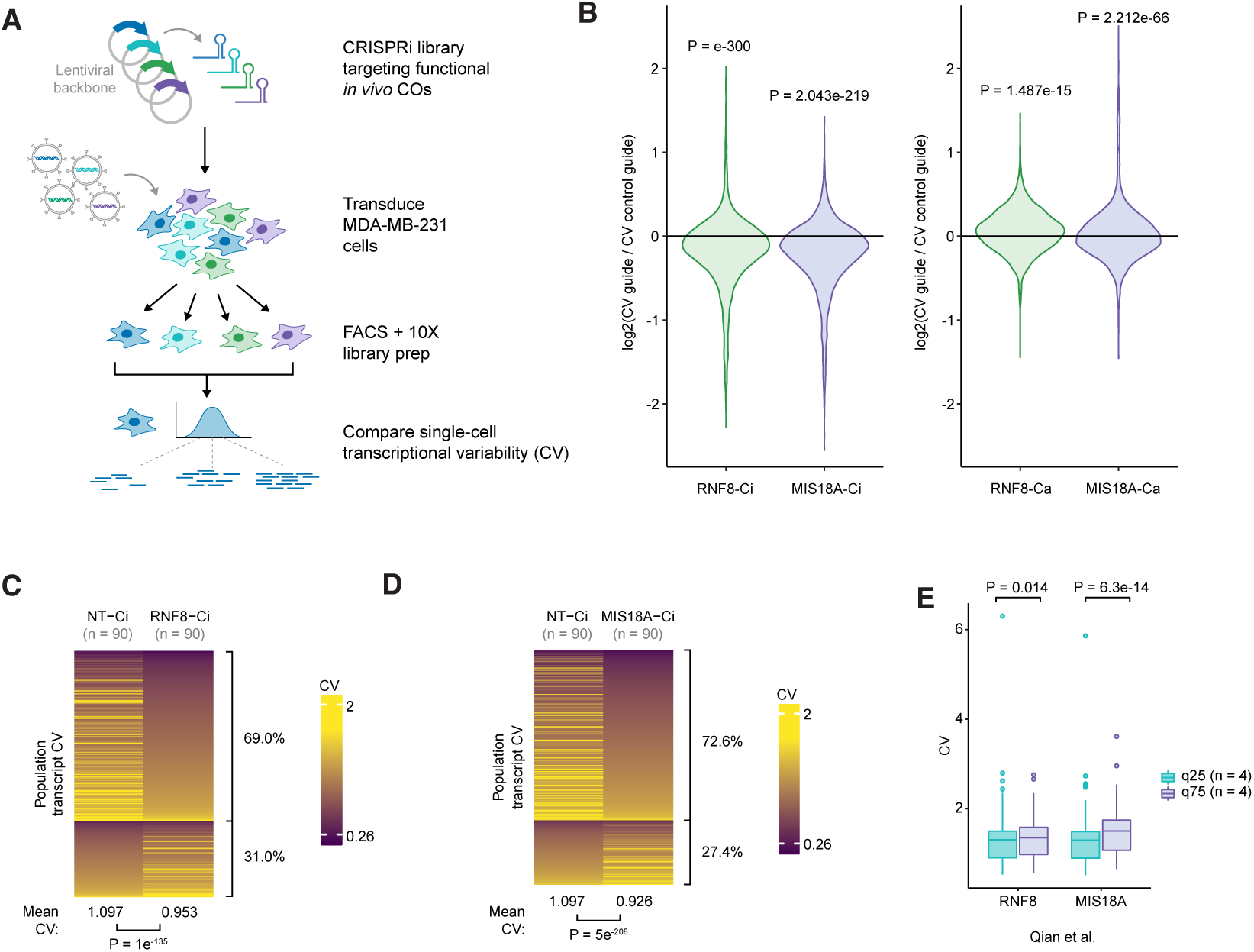
RNF8 and MIS18A modulate transcriptomic heterogeneity at the single-cell level. **(A)** Schematic of the Perturb-seq workflow. MDA-MB-231 cells were transduced with CRISPRi or CRISPRa sgRNAs targeting the 16 *in vivo*-validated COs, followed by 10X 3^′^ scRNA-seq. Genome-wide CV was computed for each perturbation and compared to size-matched random control groupings. **(B)** Mean and standard error of mean barplots for genome-wide CV change upon CRISPRi (left) or CRISPRa (right) perturbation of RNF8 and MIS18A. A value of 1 indicates no change relative to non-targeting control. *P*-values by one-sample two-tailed Student’s *t*-test (*μ* = 1). **(C, D)** Per-gene transcript CV in CRISPRi cells versus non-targeting controls for RNF8 **(C)** and MIS18A **(D)**. Each horizontal line represents a gene; *P*-values by two-sided paired Student’s *t*-test. Percentages indicate proportion of genes that decrease (top) or increase (bottom) in CV relative to non-targeting control. **(E)** CV analysis in an independent patient tumor scRNA-seq cohort ^45^. Patients were stratified into quartiles by RNF8 or MIS18A expression (*n* = 4 per quartile). *P*-values by two-sided Wilcoxon signed-rank test.

Knockdown of either RNF8 or MIS18A reduced genome-wide transcript CV relative to non-targeting controls (**Figure 3C–D**). Overexpression of either factor produced the opposite effect (**Figure S2D–E**). We validated this finding in an independent patient tumor scRNA-seq dataset ^45^, where higher RNF8 and MIS18A expression was associated with increased transcriptomic heterogeneity (**Figure 3E**). Importantly, neither RNF8 nor MIS18A expression correlated with cell cycle phase (**Figure S2F–G**), and their perturbation did not shift global mean transcript abundance (**Figure S2H–I**), indicating that these effects on transcriptomic variability are cell-cycle independent and not driven by changes in expression magnitude.

### 2.4. RNF8 and MIS18A modulate chromatin accessibility heterogeneity

RNF8 and MIS18A function in DNA damage repair and centromeric histone deposition, respectively ^36–41^, but neither has been implicated in transcriptomic heterogeneity. Given their chromatin-associated roles, we asked whether these COs modulate transcriptomic heterogeneity through changes in chromatin accessibility. We generated RNF8 and MIS18A knockdown (CRISPRi) and overexpression (CRISPRa) lines in MDA-MB-231 cells (hereafter CRISPRi/a lines) and profiled them by multiplexed scATAC-seq (**Figure 4A**; see Methods).

**Figure 4.**
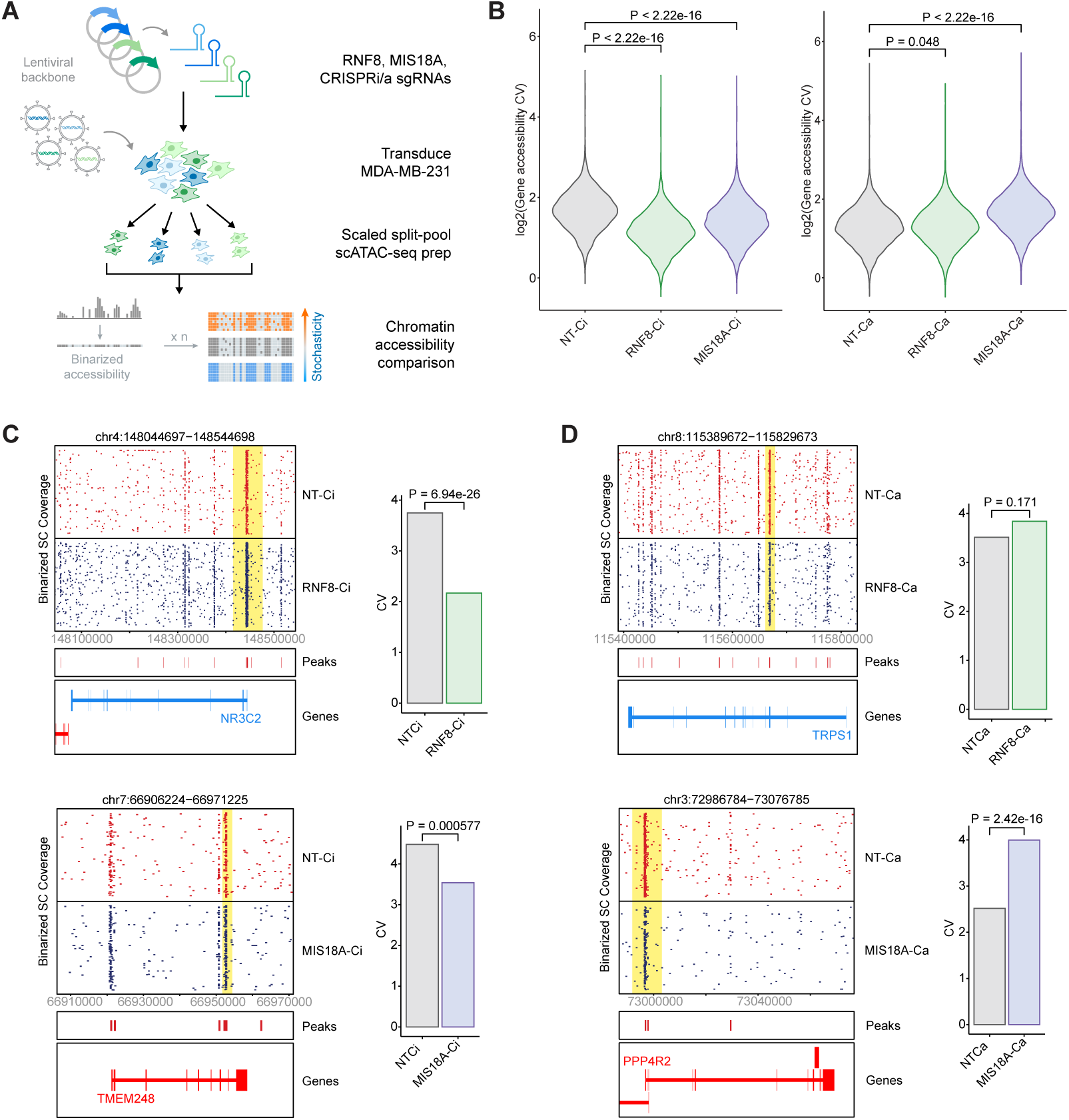
RNF8 and MIS18A expression modulates genome-wide chromatin accessibility variance. **(A)** Experimental schematic. RNF8 and MIS18A CRISPRi/a MDA-MB-231 lines were hash-pooled and pro-filed by 10X scATAC-seq. **(B)** Genome-wide open-chromatin CV change per genomic locus upon CRISPRi or CRISPRa perturbation of RNF8 and MIS18A. Accessibility was filtered for gene regions; a value of zero indicates no change relative to non-targeting control. *P*-values by two-sided Wilcoxon signed-rank test. **(C, D)** Left: binarized open/closed chromatin scores (ArchR) across cells (rows) and genomic position (columns) for representative loci. Yellow boxes highlight reduced accessibility dispersion upon knockdown **(C)** and increased dispersion upon overexpression **(D)**. Right: CV for the highlighted locus in each perturbation condition. *P*-values by Fisher’s exact test.

Genome-wide chromatin accessibility CV shifted proportionally with RNF8 or MIS18A expression (**Figure 4B**). At individual loci, knockdown of either CO drove accessibility toward uniform states (predominantly fully open or fully closed), consistent with reduced heterogeneity upstream of transcription (**Figure 4C**). Conversely, overexpression shifted loci toward intermediate open–closed ratios, indicating increased accessibility dispersion (**Figure 4D**). These changes in variance occurred without shifts in global mean accessibility (**Figure S3A–B**), indicating that RNF8 and MIS18A alter the dispersion, not the central tendency, of chromatin states. Together, these data show that the transcriptomic heterogeneity phenotypes observed by Perturb-seq are accompanied by corresponding shifts in chromatin accessibility heterogeneity.

### 2.5. MIS18A modulates transcriptomic heterogeneity through a centromere-independent pathway

MIS18A is a core component of the centromeric histone (CENP-A) deposition complex ^36–41^, raising the possibility that dysregulated centromeric heterochromatin accounts for the observed heterogeneity phenotypes. To evaluate this, we first asked whether other CENP-A deposition complex members were also associated with transcriptomic heterogeneity. HJURP, the canonical chaperone of MIS18A, scored in the TCGA bulk analysis but not in our Perturb-seq data, possibly owing to cell-cycle confounding, and MIS18B was similarly limited to the bulk modality (**Figure 1B**, **3B**, **S2A–B**). Because these results were inconclusive, we turned to more direct indicators of centromeric integrity.

CRISPRi-mediated knockdown of MIS18A reduced global CENP-A protein levels, yet CRISPRa-mediated overexpression, which increases transcriptomic heterogeneity, did not increase them (**Figure S4A**), dissociating the heterogeneity phenotype from centromeric histone levels. Loss of MIS18A can trigger accumulation of minor satellite non-coding RNAs (ncRNAs), aberrant H3K9me3/HP1 deposition, and cell death ^46–49^; however, RT-qPCR at 48 and 96 hours post-seeding revealed no significant accumulation of minor satellite ncRNAs in our CRISPRi/a lines (**Figure S4B**). Immunofluorescence for CENP-A likewise showed no change upon MIS18A knockdown (**Figure S4C**), consistent with prior reports that partial MIS18A depletion is insufficient to cause centromeric deficiency ^50^ and with our observation that CNA and aneuploidy do not explain the heterogeneity phenotypes (**Figure S1C**).

MIS18A also interacts with and is required for the activity of DNMT3A/B methyltransferases ^51^. MIS18A-directed changes in morphology heterogeneity score depended on DNMT3B activity (**Figure S4D**), suggesting that MIS18A modulates ITH through DNA methylation rather than centromeric deposition. Taken together, these data indicate that MIS18A drives transcriptomic heterogeneity and tumor progression through a centromere-independent, methylation-associated pathway.

### 2.6. Transcriptomic heterogeneity tunes chemotherapeutic sensitivity and cellular fitness

We next asked whether the transcriptomic heterogeneity driven by RNF8 and MIS18A extended to phenotypic diversity. Treatment of CRISPRi/a lines with 5-fluorouracil (5FU) or cyclophosphamide revealed that knockdown of either CO sensitized cells to both agents, whereas overexpression was protective (**Figure 5A–B**); these differences were not attributable to altered proliferation rates (**Figure S5A**). Colony formation assays showed that RNF8, but not MIS18A, perturbation altered clonogenic growth in a directionally consistent manner (**Figure 5C**), indicating that the two COs do not produce identical downstream fitness phenotypes. High-content microscopy further revealed that CRISPRi/a lines differed in morphology heterogeneity scores relative to non-targeting controls (**Figure 5D**, **S5B**; see Methods) ^50,52–55^, consistent with our earlier finding that MDA-MB-231 subpopulations with greater morphological variation exhibit increased metastatic fitness ^12^. Together, these data suggest that CO-driven transcriptomic heterogeneity contributes to phenotypic diversity and cellular fitness.

**Figure 5.**
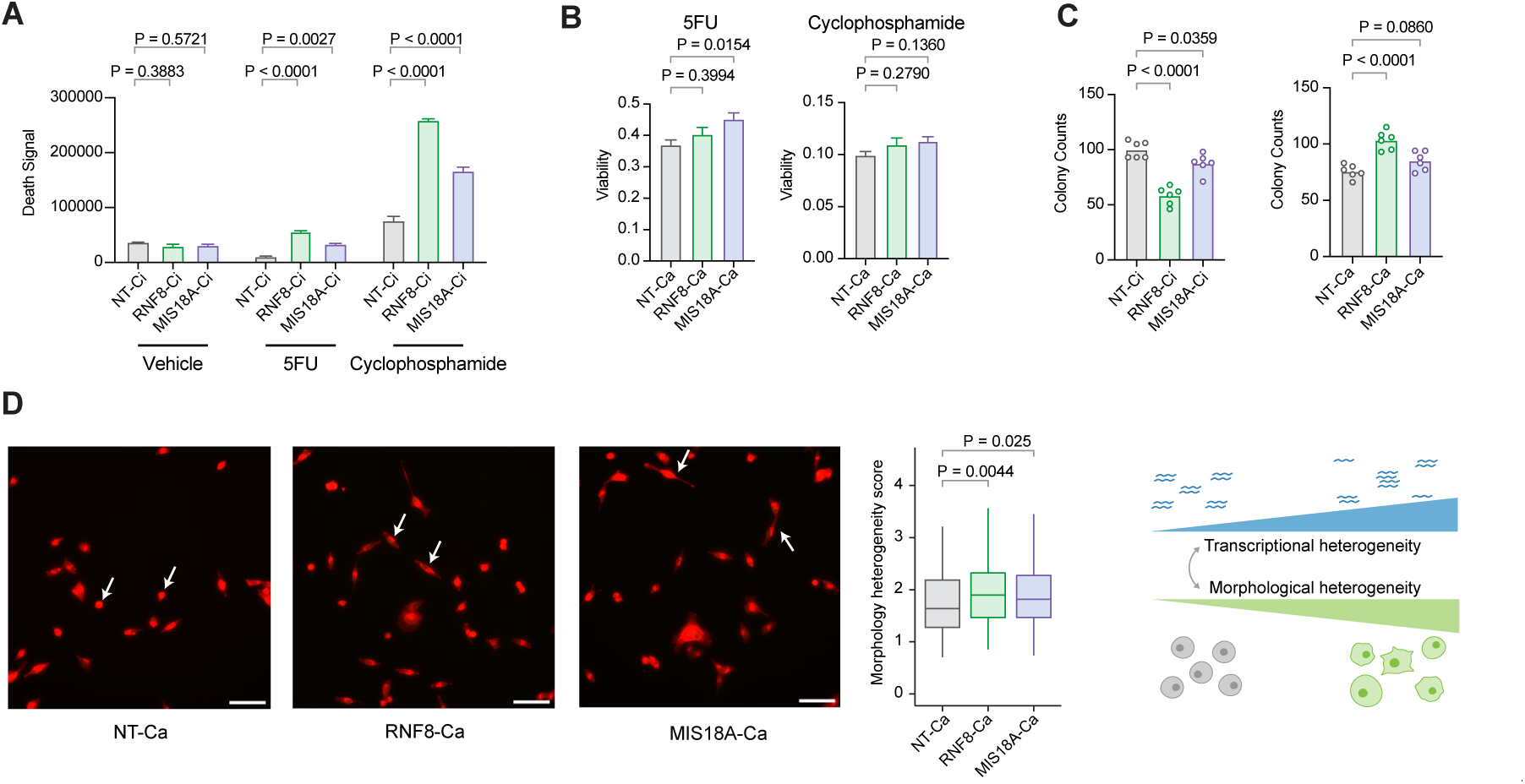
RNF8 and MIS18A tune chemotherapeutic sensitivity and proliferative fitness. **(A, B)** Cytotoxicity of MDA-MB-231 CRISPRi **(A)** and CRISPRa **(B)** lines treated with 5-fluorouracil (2.5 mM) or cyclophosphamide (2.5 mM) for 48 h. CRISPRi lines were scored by live-dead imaging (total death signal, NIRCU *μ*m^2^/image); CRISPRa lines by CellTiter-Glo 2.0 luminescence. *P*-values by one-way ANOVA (*n* = 8 per condition). **(C)** Colony formation assay of CRISPRi and CRISPRa lines transduced with the indicated sgR-NAs. *P*-values by one-way ANOVA (*n* = 6 per condition). **(D)** Cell-painting morphology heterogeneity scores for CRISPRa lines. Scores integrate cell-body, mitochondrial, ER/Golgi, and nuclear features (bar plot). Representative images shown at left; white arrows highlight morphological differences. Scale bar, 50 *μ*m. *P*-values by one-sided Wilcoxon signed-rank test.

RNF8 also functions as a post-translational stabilizer of TWIST1-directed EMT ^54^. Immunofluorescence for vimentin, a canonical EMT marker, confirmed that vimentin levels shifted upon RNF8 knockdown or overexpression (**Figure S5C**). This change did not correlate with TWIST1 transcript abundance in our scRNA-seq data (**Figure S5D**), consistent with RNF8 acting as a translational stabilizer of TWIST1 through K63 ubiquitylation ^56^.

### 2.7. RNF8 and MIS18A alter transcriptomic heterogeneity independently of gross genomic-content changes

Our bulk TCGA analysis indicated that CO-associated transcriptomic variability did not correlate with copy-number alterations (**Figure S1C**). To test this directly, we measured bulk DNA content in the CRISPRi/a lines by flow cytometry ^53–55^. Neither RNF8 nor MIS18A perturbation significantly altered DNA content in G0/G1 or G2 phases (**Figure S5E–F**), and cell-cycle phase distributions were unchanged (**Figure S5G**), consistent with our Perturb-seq observations (**Figure S2F–G**). DNA content CV across phases was also unaffected (**Figure S5H**). Because our TCGA analysis used SNP-array CNA data and our cell-line measurements used flow cytometry, both chromosomal and extrachromosomal CNA events ^57^ are captured by these assays. Together, these results indicate that RNF8 and MIS18A drive transcriptomic heterogeneity through mechanisms independent of CNA and aneuploidy.

### 2.8. RNF8 and MIS18A drive tumor progression and predict patient survival

To link transcriptomic heterogeneity to metastatic progression, we injected our MDA-MB-231 CRISPRi/a lines into mice and monitored lung colonization. Knockdown of either RNF8 or MIS18A reduced metastatic burden, whereas overexpression increased it (**Figure 6A–B**). This phenotype was not cell-type-specific: an independently engineered HCC1806 line recapitulated the result (**Figure S6A**).

**Figure 6.**
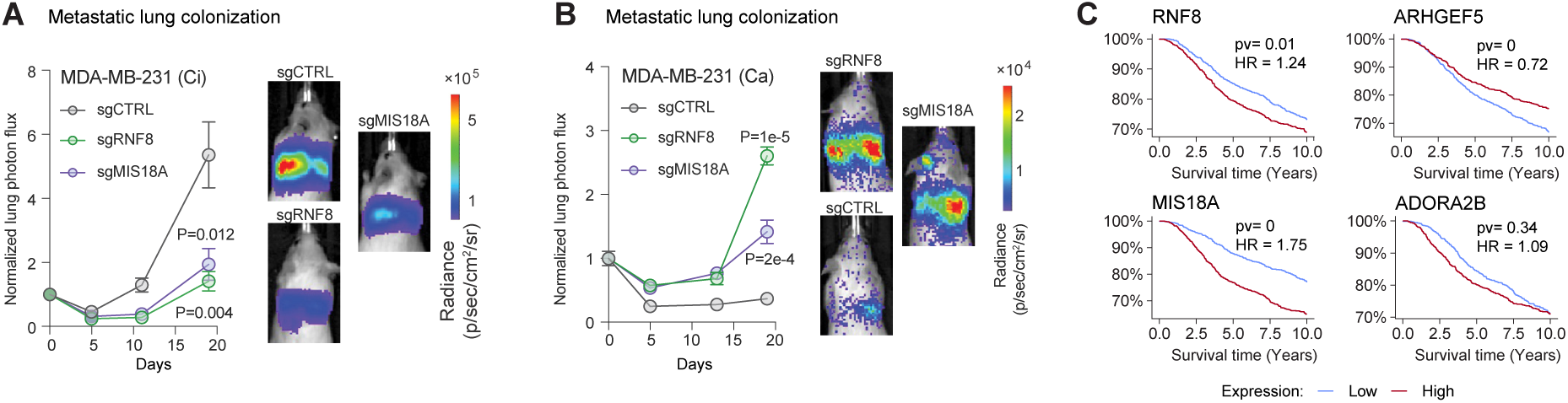
RNF8 and MIS18A drive metastatic progression *in vivo* and predict patient survival. **(A, B)** Lung metastatic burden (normalized photon flux) over time for MDA-MB-231 CRISPRi **(A)** and CRISPRa **(B)** lines. *P*-values by one-tailed *t*-test (*n* = 5 per sgRNA condition). **(C)** Kaplan–Meier survival curves for METABRIC patients stratified into bottom-quartile (red) and top-quartile (blue) expressors of the indicated chromatin organizers. Control genes shown at right. *P*-values by log-rank test.

We next examined clinical relevance using the METABRIC dataset ^58^. Higher expression of RNF8 or MIS18A was associated with shorter patient survival, and lower expression with longer survival (**Figure 6C**). A similar trend was observed across other CENP-A deposition complex members (**Figure S6B**) ^36,41^. Because these survival associations span several complex components, yet are not uniformly shared across all of them, they likely reflect broader clinical relevance of centromere/kinetochore-linked programs (e.g., proliferation, aneuploidy propensity) rather than direct evidence that the entire complex modulates ITH. Among these components, MIS18A showed the most reproducible heterogeneity phenotype in our single-cell perturbation assays, motivating the focused mechanistic dissection described above. Notably, CENP-A itself was preferentially associated with metastatic over primary tumor progression in our *in vivo* CRISPRi data (**Figure S6C**). Together, these results establish RNF8 and MIS18A as drivers of transcriptomic heterogeneity that promote metastatic progression *in vivo* and predict clinical outcome in human breast cancer.

## 3. Discussion

Intratumoral transcriptional heterogeneity (ITH) is increasingly recognized as a clinically important feature of aggressive tumors ^1,6–8^, yet strategies for systematically identifying non-genetic regulators of this phenotype have remained limited. Here, we present a framework for discovering candidate modulators of ITH by integrating bulk patient transcriptomic data, functional *in vivo* screening, and single-cell transcriptomic and epigenomic readouts. Applying this framework to breast cancer, we identify RNF8 and MIS18A as chromatin-associated factors whose perturbation produces coordinated shifts in transcriptomic variability, chromatin accessibility heterogeneity, and cancer-relevant phenotypes including chemotherapeutic resistance, morphological diversity, and metastatic progression *in vivo*. More broadly, the approach provides a generalizable template for prioritizing candidate regulators of ITH in other tumor types and disease contexts.

Prior work in yeast and, more recently, mammalian cells has established that specific genes can act as “stochastic tuners,” modulating the breadth of transcriptomic states accessible to isogenic populations ^13,35^. Our findings extend this concept to a cancer setting by identifying chromatin organizers that function as endogenous modulators of transcriptomic dispersion. RNF8 and MIS18A were nominated through multiple orthogonal screening filters and show directionally consistent effects across single-cell transcriptomic and chromatin accessibility variability. These heterogeneity phenotypes were not accompanied by global shifts in mean transcript abundance, cell-cycle redistribution, or bulk DNA-content changes, supporting the interpretation that RNF8 and MIS18A influence ITH through non-genetic, chromatin-level mechanisms in our tested models. While our data do not directly establish long-term epigenetic inheritance, the concordance between scRNA-seq and scATAC-seq variability supports a model in which these COs alter the range of accessible regulatory states that cells can sample.

RNF8 is best characterized for its role in DNA damage response (DDR) signaling and for post-translational regulation of EMT-related programs through K63-mediated ubiquitylation of TWIST1^40,54,56^. Both functions provide plausible routes by which RNF8 activity could influence chromatin remodeling and downstream transcriptional-state diversity. DNA double-strand break repair can transiently reshape local chromatin architecture and, in some contexts, leave durable epigenetic changes after repair ^59,60^. We do not directly assess damage-induced chromatin remodeling here, but DDR pathway engagement and repair-pathway choice represent candidate mechanisms by which RNF8 could expand chromatin-state variability during tumor evolution.

MIS18A is canonically linked to centromeric chromatin and CENP-A deposition ^36,41^, yet several lines of evidence argue that the heterogeneity phenotypes we observe operate through a centromere-independent pathway. MIS18A overexpression, which increases transcriptomic heterogeneity, did not increase CENP-A protein levels; minor satellite ncRNA accumulation and CENP-A localization were unaffected; and gross DNA-content changes were absent. Instead, MIS18A-directed changes in morphology heterogeneity depended on DNMT3B activity, pointing to DNA methylation as the likely effector pathway ^51^. MIS18A is a known interaction partner and activator of DNMT3A/B methyltransferases, and perturbation of DNA methylation patterns could plausibly alter the stability and reversibility of chromatin states across a cell population. Characterizing the specific genomic targets of MIS18A-dependent methylation changes will be an important goal for future mechanistic studies.

Our observation that knockdown of RNF8 or MIS18A sensitized cells to 5-fluorouracil and cyclophosphamide, while overexpression conferred resistance, provides direct functional evidence linking transcriptomic heterogeneity to therapeutic response. This is consistent with the hypothesis that transcriptional plasticity enables cancer populations to access a wider distribution of phenotypic states under selective pressure, increasing the probability that resistant subpopulations pre-exist at the time of treatment ^13,35^. From a translational perspective, these findings raise the possibility that targeting variance-promoting chromatin regulators could narrow the phenotypic landscape available to tumor cells and thereby enhance chemotherapeutic efficacy, particularly as a combination strategy alongside conventional cytotoxic agents.

Overall, these data establish that modulators of transcriptional variability represent a distinct and actionable axis of tumor biology. A practical contribution of this work is the demonstration that combining bulk patient datasets with functional screening and single-cell readouts can systematically prioritize candidate “variance regulators” for mechanistic study. We anticipate that applying this framework across additional tumor types and extending it to other classes of non-genetic regulators will reveal further targets whose inhibition could constrain the phenotypic diversity that fuels drug resistance and metastatic progression.

## 4. Methods

### 4.1. Experimental Model and Study Participant Details

#### 4.1.1. Cell lines and Cell culture

MDA-MB-231 breast cancer cell line was acquired from ATCC. All cells were cultured in a 37^◦^C 5% CO_2_ humidified incubator. MDA-MB-231 was cultured in DMEM medium supplemented with 10% FBS, glucose (2 g/L), L-glutamine (2 mM), 25 mM HEPES, penicillin (100 units/mL), streptomycin (100 *μ*g/mL) and amphotericin B (1 *μ*g/mL) (Gibco). All cell lines were routinely screened for mycoplasma with a PCR-based assay. To select transgenic lines, puromycin was used at 8 *μ*g/mL final concentration.

#### 4.1.2. Mouse Models

Female NSG mice were purchased from Jackson Laboratory (Strain#005557). All animal surgeries, husbandry and handling protocols were completed according to University of California IACUC guidelines.

### 4.2. In vivo CRISPR screening

#### 4.2.1. In silico nominated sgRNA library cloning and sequencing validation

For our computationally nominated CRISPRi library, a library consisting of guides targeting 185 elements (5 sgRNAs each for 37 genes) was designed and ordered from Twist Biosciences. The pool was resuspended to 5ng/*μ*L final concentration in Tris-HCl 10mM pH 8, and a qPCR to determine Ct to be used for down-stream library amplification was performed (forward primer: TCACAACTACACCAGAAGccac, reverse primer: TCTTCGTCAAAGTGTTGCcagc) using a 16-fold library dilution.

The library was then amplified via PCR, and ran out on a 2% agarose gel to check library size (expected band of 84bp). PCR product was then cleaned up using a DNA Clean and Concentrator kit-5 (Zymo Research Cat. #D4003), and eluted in 15*μ*L H_2_O. Cleaned product was digested overnight using FD Bpu1102I (Thermo Fisher Cat. #FD0094), and then further digested for 1hr using FD BstXI (Thermo Fisher Cat. #FD1024). Inserts were then ligated into pCRISPRi/a v2 backbone in a 50ng reaction with 1:1 insert:backbone ratio for 16hrs 16C. Ligated products were then ethanol-precipitated overnight at −20C, cleaned, and then transformed into 100*μ*L NEB Stables (NEB Cat. #C3040H), followed by maxiprep plasmid isolation.

For sequencing validation, 1*μ*g plasmid DNA was then digested in 50*μ*L volume for 1hr with FD BstXI (Thermo Fisher Cat. #FD1024). Digested plasmid DNA was then Klenow-extended using added UMI linker (sequence: CTCTTTCCCTACACGACGCTCTTCCGATCTNNNNNNcttg), and then cleaned up using a Zymo DNA Clean & Concentrator-25 kit (Zymo Research Cat. #D4033). Indexing PCR (forward primer: AATGATACGGC-GACCACCGAGATCTacactctttccctacacgacgctc; reverse primer: CAAGCAGAAGACGGCATACGAGATGATCTGGT-GACTGGAGTTCAGACGTGTGCTCTTCCGATcgactcggtgccactttttc) was then performed in 30*μ*L final volume, followed by gel purification (Takara Bio Cat. #740609.50). Samples were then pooled and sequenced on a lane of HiSeq 4000 SE ^47^ at the UCSF Center for Advanced Technology (CAT).

#### 4.2.2. Viral titering of computationally nominated sgRNA library

3 million HEK293Ts were seeded in a 10cm plate. 24hrs later, HEK293Ts were transfected with TransIT-Lenti (Mirus Bio Cat. #Mir6603) reagent according to manufacturer’s protocol. Viral supernatant was harvested, aliquoted, flash-frozen, and then stored −80C for long-term storage.

200K MDA-MB-231 CRISPRi-ready cells were then seeded in a 6-well plate for viral titering. Using a range of 100-, 200-, and 400*μ*L viral supernatant, cells were transduced, adding polybrene to 8 *μ*g/mL final concentration. 48hrs post-transduction, cells were passed through flow cytometry on the FACS Aria II in the UCSF CAT, and %BFP+ was recorded.

#### 4.2.3. Cell preparation for subcutaneous injection

10 million MDA-MB-231 CRISPRi-ready cells were seeded into a 15cm plate and grown overnight. On the following day, lentivirus was added to cells with a target MOI of <10%, with polybrene added to final concentration 8 *μ*g/mL. Media was then changed 24hrs post-transduction, and selection was started at 72 hrs post-transduction via puromycin at final concentration 1.5 *μ*g/mL.

We then partitioned into 3 arms the transduced MDA-MB-231 CRISPRi-ready cells. Specifically, 200K cells were split into a 15cm plate for in vitro passage for sgRNA in vitro growth normalization. 200K cells were pelleted and frozen at −80C for downstream gDNA extraction, for ‘t0’ collection.

For the bilateral flank injections, 9 million cells were spun down and resuspended to final concentration 1 million cells/50*μ*L in 1:1 PBS/matrigel. Bilateral subcutaneous injections in 50*μ*L final volume were then performed in female, 8-12 week-old age-matched NOD scid gamma (NSG) mice (n = 3) with 500K cells.

For lung colonization assay, 1 million cells were spun down and resuspended to final concentration 100K/100 *μ*L in PBS. Tail-vein injections in 100*μ*L final volume were then performed in female, 8-12 week-old age-matched NOD scid gamma (NSG) mice.

#### 4.2.4. Tumor gDNA extraction and library preparation

Tumors were then harvested 5-6 weeks post-injection and gDNA extracted using Quick-DNA midiprep plus kit (Zymo Research Cat. #D4075).

For the bilateral flank injections, each tumor gDNA sample (n = 6) was digested in 3 *μ*g-scale, 100*μ*L volume reactions with FD BstXI. Digested gDNA was then Klenow-extended using added UMI linker (se-quence: CTCTTTCCCTACACGACGCTCTTCCGATCTNNNNNNcttg), and then cleaned up using a Zymo DNA Clean & Concentrator-25 kit (Zymo Research Cat. #D4033), eluting twice in 50 *μ*L Qiagen EB pre-heated to 70C (final elution volume of 100 *μ*L). Indexing PCRs (forward primer: AATGATACGGCGACCACCGAGATCTacactctttccctacacgacgctc; reverse primer: CAAGCAGAAGACGGCATACGAGATGATCTGGTGACTGGAGTTCA-GACGTGTGCTCTTCCGATcgactcggtgccactttttc) were then performed with 500ng tagged gDNA in 100*μ*L final volume with the following parameters: 98C 30s, [98C 30s 62C 15s 72C 15s 30X], 10C hold, followed by 150-1000bp double-sided cleanup (Zymo Research Cat. #D4085). Samples were then pooled and sequenced on a lane of HiSeq 4000 SE ^47^ at the UCSF Center for Advanced Technology (CAT).

For harvested lungs from lung colonization assay (n=5), lungs were first mouse-cell-depleted using Miltenyi Mouse Cell Depletion kit (Miltenyi Cat. # 130-104-694) to enrich human MDA-MB-231 cell line xenograft signal. Tumor gDNA was then digested in 3 *μ*g-scale, 100 *μ*L volume reactions with FD BstXI. Digested gDNA was then Klenow-extended using added UMI linker (sequence: CTCTTTCCCTACACGACGCTCTTCCGATCTNNNNNNcttg), and then cleaned up using a Zymo DNA Clean & Concentrator-25 kit (Zymo Research Cat. #D4033), eluting twice in 50 *μ*L Qiagen EB pre-heated to 70C (final elution volume of 100 *μ*L). Indexing PCRs (forward primer: AATGATACGGCGACCACCGAGATCTacactctttccctacacgacgctc; reverse primer: CAAGCAGAA-GACGGCATACGAGATGATCTGGTGACTGGAGTTCAGACGTGTGCTCTTCCGATcgactcggtgccactttttc) were then performed with 500ng tagged gDNA in 100*μ*L final volume with the following parameters: 98C 30s, [98C 30s 62C 15s 72C 15s 30X], 10C hold, followed by 150-1000bp double-sided cleanup (Zymo Research Cat. #D4085). Samples were then pooled and sequenced on a lane of HiSeq 4000 SE ^47^ at the UCSF Center for Advanced Technology (CAT).

### 4.3. In vitro scRNA-seq

#### 4.3.1. scRNA-seq sgRNA library cloning

For our CRISPRi library, a library consisting of guides targeting a total of 16 elements, consisting of COs with |z| > 4 across both MFP and TV screens, along with non-targeting sgRNAs, was designed and ordered from Twist Biosciences. We selected the top 2 predicted in silico protospacers and ordered them as a paired sgRNA cassette for cloning into a compatible dual sgRNA lentiviral CRISPR guide vector, pJR85 (Addgene Cat. #140095). The pool was resuspended to 10ng/*μ*L final concentration in Tris-HCl 10mM pH 8, and an initial PCR to amplify the oligo pool (forward primer: TCACAACTACACCAGAAGccac, reverse primer: TCTTCGT-CAAAGTGTTGCcagc) was performed with the following cycling conditions: 98C 30s, [98C 15s, 56C 15s, 72C 15s 11X], 72C 1 min, 10C hold. The library was then purified via Qiagen Min Elute kit (Qiagen Cat.# 28004), eluted in 20 *μ*L Qiagen EB, and ∼0.8ug was recovered post-elution. Purified insert was then digested overnight at 37C with BstXI (Thermo Fisher Cat. #FD1024) and Bpu1102l (Thermo Fisher Cat. #FD0094), and run out on a 8% TBE gel. The expected insert size of 97bp was cut and extracted via ethanol precipitation (using 3X volume 100% EtOH) overnight at −20C. Purified insert was then ligated into BstXI/Bpu1102l-digested pJR85 at a 1:1 molar ratio for 16 hrs 16C, and ligation product was purified via ethanol precipitation overnight at −20C. Final precipitation product was eluted in 5 *μ*L H_2_O and used as input to electroporation using 50 *μ*L MegaX electrocompetent cells (Invitrogen Cat. #C640003), and culture was prepped in a maxiprep format.

For the second ligation, a golden gate assembly reaction was set up with pJR89 donor vector and the generated pJR85 intermediate library described above, with the following cycling conditions: [42C 5mins, 16C 5 mins 30X], 60C 5 mins. Product was ethanol-precipitated at −20C, resuspended in 10 *μ*L H_2_O, and transformed into 100 *μ*L NEB Stable (New England Biolabs Cat. #C3040H). 50mL of resulting library transformants was used for midiprep plasmid isolation.

For sequencing validation, 200ng of plasmid library was used for PCR (forward primer: AATGATACG-GCGACCACCGAGATCTACACCGCGGTCTGTATCCCTTGGAGAACCACCT, reverse primer: CAAGCAGAAGACG-GCATACGAGATcgtgaGCGGCCGGCTGTTTCCAGCTTAGCTCTTAAA) with the following cycling conditions: 98C 30s, [67C 10s, 72C 75s 12X], 72C 5 mins, 10C hold. The product was size selected for >150bp fragments and quantified via tapestation, and the library was sequenced using a MiSeq v2 kit PE150 configuration.

#### 4.3.2. Viral titering

0.5 million HEK293Ts were seeded in a 10cm plate. 24hrs later, HEK293Ts were transfected with the plasmid library using TransIT-Lenti (Mirus Bio Cat. #Mir6603) reagent. Viral supernatant was harvested, aliquoted, flash-frozen, and then stored −80C for long-term storage.

200K MDA-MB-231 CRISPRi-ready cells were then seeded in a 6-well plate for viral titering. Using a range consisting of 250 *μ*L, 500 *μ*L, and 1mL viral supernatant, cells were transduced with polybrene at 8 *μ*g/mL final concentration. 24h post-transduction, media was replaced with fresh media without polybrene. 48hrs post-transduction, cells were passed through flow cytometry on the FACS Aria II in the UCSF CAT, and %BFP+ was recorded for each condition, and was used to record viral titer for the frozen virus library.

#### 4.3.3. scRNA-seq library workflow & sequencing

1 million MDA-MB-231 CRISPRi-ready cells were seeded in a 15cm plate. 24h after seeding, 2mL of virus was added to the plate with polybrene at 8 *μ*g/mL final concentration. 24h after transduction, the media was changed to polybrene-free media. 48h after transduction, cells were trypsinized and passed on FACS, and BFP+ cells were isolated to be used as input to 10X scRNA-seq. ∼100K BFP+ cells were isolated from flow, spun down at 800g 5 mins RT, and resuspended in 100 *μ*L PBS. Cells were loaded at a target of 10K cells on a single lane of 10X scRNA-seq v3.1, and the manufacturer’s protocol was followed for library preparation. The resulting indexed library was sequenced on a lane of NovaSeq 6000.

### 4.4. Colony formation assay

For colony formation assay, 200 cells per relevant MDA-MB-231 cell line were seeded (n=6) in a 6-well plate. 8 days after seeding, colonies were stained and imaged. Briefly, media was removed and cells were washed with 1mL PBS at RT. Cells were then fixed in 4% PFA (Alfa Aesar Cat. #43368-9L) for 10 minutes at RT, and then stained in 0.1% crystal violet (Sigma-Aldrich Cat. #V5265-250ML) for 1h at RT. Wells were then washed 3X with ddH_2_O at RT until colonies were visible. Colonies were then imaged on a Bio-Rad ChemiDoc MP imager.

### 4.5. Cell proliferation and cytotoxicity assays

For assaying cytotoxicity, two assays were used. For the CRISPRa data shown in the main text, CellTiter-Glo 2.0 Cell Viability Assay (Promega Cat. #G9241) was used. 5K of the relevant MDA-MB-231 cell line was seeded per well in a black 96-well plate (Corning Cat. #3904) for luminescence measurement. 8 wells were seeded per cell condition in 100*μ*L volume media. 24h after seeding, cell media was replaced with media containing either 5FU or cyclophosphamide at [5FU]: 5mM; [cyclophosphamide]: 2.5mM. 48h after drug treatment, cells were harvested according to manufacturer’s protocol. Briefly, CellTiter-Glo 2.0 Reagent and cell plates were equilibrated to RT 30 mins prior to use. 100*μ*L CellTiter-Glo 2.0 Reagent was then added via multichannel to each well and mixed at 300 rpm for 2 mins at RT; the plate was incubated for 10 minutes at RT, covered. Plate luminescence was then recorded on a Tecan Spark Microplate reader.

For the CRISPRi data in the main text, cells were imaged in real time using an Incucyte SX5. 5K of the relevant MDA-MB-231 cell line was seeded per well in a black 96-well plate (Corning Cat. #3904). 8 wells were seeded per cell condition in 100*μ*L volume media. 24h after seeding, cell media was replaced with media containing 1X NIR live-dead dye (Sartorius Cat. #4846) along with either 5FU or cyclophosphamide at [5FU]: 10mM; [cyclophosphamide]: 2.5mM. The plate was placed in the Incucyte and imaged at 3-hour intervals at 4X objective with 3 images per image snapshot. Death signal was recorded via the respective NIR Incucyte channel, and total integrated intensity (NIRCU × *μ*m^2^/image) was used in the comparison between MDA-MB-231 cell lines with respective treatments.

### 4.6. High-content microscopy

The JUMP v3 kit (Revvity Cat. # PING21) was purchased and used to carry out Cell Painting morphological assaying ^59^. Briefly, 5K of each respective MDA-MB-231 CRISPRi or MDA-MB-231 CRISPRa line were seeded in replicates in a black 96-well plate (Corning Cat. #3904). 24h after seeding, 50 *μ*L of staining solution 1 was added to each well and placed in an incubator at 37C, 5% CO_2_ for 30 mins. Cells were then fixed with 50 *μ*L 16% PFA (Alfa Aesar Cat. #43368-9L) for 20 mins at RT in the dark. For cell feature staining, 50 *μ*L of staining solution 2 was added to each well for 30 mins at RT in the dark. The plate was then stored at 4C in the dark in 0.3% NaN3 solution until image processing.

### 4.7. In vivo tail vein injections for individual hit validation

For in vivo lung colonization assay, MDA-MB-231 (CRISPRi-ready, or CRISPRa-ready with appropriate sgRNA) were grown in 10cm plates and allowed to expand. On the day of injections, cells were harvested and resuspended to final concentration 200K/100 *μ*L in PBS. Tail-vein injections in 100 *μ*L final volume were then performed in female, 8-12 week-old age-matched NOD scid gamma (NSG) mice (Jackson labs). In vivo bioluminescence was monitored weekly by (intraperitoneal) injection of luciferin and normalized to bioluminescence signal immediately following cell injection.

### 4.8. scATAC-seq of individually generated lines

#### 4.8.1. Nuclei extraction

To isolate nuclei from cells as input to scATAC-seq, cells were trypsinized and 5 x 10^6^ cells were spun at 300xg for 5 mins at 4C. Supernatant was removed and cells were then lysed in 1mL lysis buffer (10mM Tris-HCl pH 7.4, 10mM NaCl, 3mM MgCl2, .03% IGEPAL-630 (Millipore-Sigma Cat. #I8896-50ML)) for a total of 15 minutes on ice. Cells were then washed 2X in 1mL wash buffer (1% BSA/PBS), spun down at 500xg for 10 mins at 4C, and resuspended in 1X 10X Nuclei buffer (10X Genomics Cat. #2000207). Nuclei isolation was confirmed via trypan blue staining and live-dead quantification on a Countess III FL. For the scATAC-seq experiment described in the main text, >99% death was observed, corresponding to isolated nuclei.

#### 4.8.2. Scale pre-indexing and cell line hashing

For higher throughput and to address batch effects that would arise from loading separate samples on separate channels on the 10X platform, nuclei that were isolated were then used as input to the Scale pre-indexing kit for scATAC-seq (Scale Cat. #110001), and the manufacturer’s protocol for hashing was followed. Briefly, nuclei for each of the 6 MDA-MB-231 cell lines were diluted to a loading concentration of 4K nuclei/uL (for a target of 20,000 nuclei/well). 5 *μ*L was loaded per well of the provided ITP, and 4 wells were loaded for each of the 6 MDA-MB-231 cell lines. Nuclei were pooled and diluted to a final concentration of 7.142K/uL in the provided loading buffer, and 100K nuclei were loaded per channel, 2 channels total, of a 10X Chromium X controller.

#### 4.8.3. scATAC-seq of isolated MDA-MB-231 nuclei

For scATAC-seq library generation, we used the 10X scATAC-seq library v2 kit (10X Genomics, PN-1000390). The protocol was followed as described starting from step 2 of the protocol, with the following modifications: in step 2.5a, 4 cycles were used; steps 3.1p, 3.2a/k, brief vortexing was used to mix instead of pipette mixing; step 3.2l, a 5 minute RT incubation was used; step 4.1c, 2.5 *μ*L of either is701, is702 primer (Scale Cat. #110101) was used instead of 10X single Index N Set A primer; step 4.1d, 8 cycles were used; step 4.2a/e/n, brief vortexing was used to mix instead of pipette mixing; step 4.2o: 5 mins RT incubation was used.

### 4.9. DNA content analysis of generated MDA-MB-231 CRISPRi/a lines

1 million cells per cell line were trypsinized and spun down for 5 mins 500xg at RT. Human PBMCs were also included during this process to serve as internal DNA standard ^46–48^. Cells were resuspended in ∼0.5mL PBS, and then 70% ice-cold EtOH was added dropwise to cells over the course of ∼30s with constant vortexing. Cells were then spun down for 10 mins at 500xg and cells were washed 1X with PBS. Post-wash, cells were resuspended in 1mL PI staining buffer (40 *μ*g/mL PI (Sigma-Aldrich Cat. #P4864-10ML); 100 *μ*g/mL RNAse A (Thermo Fisher Cat. #EN0531)) for 1 hr at RT in the dark before running on flow cytometry.

### 4.10. Immunoblotting and Immunohistochemistry

#### 4.10.1. Cell lysates

For immunoblotting, cells were seeded in 6-well plates and harvested at confluency. Cells were washed 3X in 1X PBS, and then lysed in 200 *μ*L 1X RIPA buffer with supplemented protease inhibitor (50mM Tris pH 8.0, 150mM NaCl, 1% IGEPAL CA-630, 1% sodium deoxycholate, 0.1% SDS) for 10 mins on ice. Cell lysates were then passed through a 28g needle 2X to shear gDNA, spun down at max speed 4C, and stored at −80C for long-term storage.

#### 4.10.2. BCA protein quantification assay

For BCA assay, working solution was performed according to manufacturer’s protocol. Briefly, working reagent was made by mixing reagent B:A in a 50:1 ratio. Samples including standards were incubated in 200 *μ*L volume at 37C for 30 mins and allowed to cool to RT. Samples were then read on a nanodrop and sample concentrations were recorded.

#### 4.10.3. Gel running

Samples were run on a NuPage 4-12% gradient gel in MOPS SDS running buffer. 15ug of protein were loaded per well in 20 *μ*L volume at 200V for 50 mins. For transfer, the iBlot3 system was used, and the resulting transfer membrane was checked with Ponceau S staining solution for proper transfer.

#### 4.10.4. Blocking and antibody incubation

Membrane was incubated in blocking buffer (5% non-fat milk/PBST). After block, primary antibody was added to each cut membrane portion at appropriate dilution in antibody staining buffer (2% BSA/PBST) and incubated overnight on a rocker at room temperature. Membrane was then washed 3X in 1X PBST for 10 mins on a rocker. Secondary antibody was then added at 1:10,000 dilution in antibody staining buffer (2% BSA/PBST) and then incubated for 1hr on a shaker with aluminum foil. Membrane was then washed 3X in 1X PBST for 10 mins with foil on, and then imaged on an Odyssey fluorescence imager.

### 4.11. Minor satellite ncRNA qRT-PCR

6-well plates were seeded with MDA-MB-231 CRISPRi/a cell lines at 300K cells/well (n=4 per condition). Cells were allowed to grow for 48hrs or 96hrs to allow for accumulation of minor satellite ncRNA species, and RNA was then extracted from each well using a Qiagen microprep RNA kit. cDNA was then constructed from each RNA sample (RT mixture: 300ng RNA, 5X RT buffer, 6.25ng random hexamers, 0.125 *μ*L oligo dT 100uM, 0.25 *μ*L dNTP 10mM, 0.125 *μ*L RNAseOUT, .0625 *μ*L Maxima H Minus RT, 200u/ul, 0.3125 *μ*L ddH_2_O) with the following thermal incubation parameters: [10mins 25C, 15 mins 50C, 5mins 85C]. cDNA was then diluted 10-fold in ddH_2_O to a final volume of 50 *μ*L, added to qPCR master mix solution (5 *μ*L 2X qPCR MM, 0.3 *μ*L primer F 10uM, 0.3 *μ*L primer R 10uM, 2 *μ*L cDNA, 2.4 *μ*L ddH_2_O), and cycled with the following parameters on a Roche Lightcycler 480: 2 mins 95C, [15s 95C, 45s 60C] x 35.

### 4.12. EMT immunofluorescence

Cells were seeded at a density of 5×103 cells per well in a 96-well plate in culture medium and incubated for 72 hours. Wells were rinsed three times with PBS-T, and cells were fixed with 5% paraformaldehyde (pH 7.4) for 15 minutes at room temperature, followed by three washes with PBS-T. Cells were permeabilized with 0.1% Triton X-100 in PBS for 5 minutes at room temperature and washed three additional times. Blocking was performed with 10% goat serum (Thermo Fisher #50197Z) in PBS for 60 minutes at room temperature.

For vimentin staining, rabbit anti-vimentin antibody (Abcam #ab92547, 2 *μ*g/mL) was diluted in 1% goat serum and incubated overnight at 4^◦^C. After washing, goat anti-rabbit Alexa Fluor 488 highly cross-adsorbed secondary antibody (Thermo Fisher #A-11034, 2 *μ*g/mL) was diluted in 0.1% goat serum and incubated for 45 minutes at room temperature protected from light. Secondary-only controls were included for each stain. After final washes, plates were imaged using a Molecular Devices ImageXpress high-content imaging system.

### 4.13. Cell painting assay

Cells were seeded at a density of 5×103 cells per well in 96-well plates in culture medium containing compound or vehicle. Nanaomycin A and DY-46-2 were each tested at 11 doses in a 2-fold serial dilution series with a top concentration of 5 *μ*M, plus a DMSO vehicle control. Cells were incubated at 37^◦^C, 5% CO_2_ for 72 hours. Cell painting was performed using the PhenoVue Cell Painting JUMP Kit (Revvity, #PING21) according to the manufacturer’s protocol and imaged using a Molecular Devices ImageXpress high-content imaging system.

### 4.14. Image analysis

Immunofluorescence images were analyzed in FIJI with the following parameters: DAPI = 200ms, TRITC (CENP-A) = 280ms, trans = 15ms. For nuclear quantification, the DAPI channel was background-adjusted using auto min/max display scaling and smoothed with a median filter. Nuclei were then identified using a local-maxima–based detection workflow with a fixed prominence threshold, and counts were recorded per field. The corresponding TRITC channel was processed using the same general settings to enable comparison across conditions. Identical analysis parameters were applied within an experiment whenever possible, with obviously oversaturated images noted during quality control.

Vimentin-stained images were quantified in Fiji/ImageJ using a mask-based intensity workflow. Images were converted to 32-bit and background corrected using rolling-ball subtraction. A duplicate image was smoothed and thresholded to generate a binary signal mask, which was refined by simple morphological cleanup. Signal intensity was then measured on the background-corrected original image within the mask, while background intensity was estimated from the inverse masked region. Background-corrected summary metrics, including median intensity, mean intensity, and integrated density–based measurements, were exported for downstream comparison across conditions.

### 4.15. Immunohistochemistry

6-well plates were seeded with MDA-MB-231 CRISPRi/a cell lines at 50K cells/well and allowed to grow for 24hrs prior to staining. Wells were covered to 3mm depth with 4% formaldehyde and allowed to sit for 15 mins at room temperature, then washed 3X with 1X PBS. Permeabilization buffer (0.2% IGEPAL-630 in PBS) was then added to cells for 5 mins at room temperature, and washed 3X with 1X PBS. Cells were then blocked with buffer (2% BSA/PBS) for 60 mins at room temperature, and appropriate primary antibody was applied overnight at 4C (anti-CENPA, Invitrogen Cat. #MA1-20832). Plate was then washed 3X with 1X PBS, and incubated with appropriate secondary antibody for 1hr at room temperature (Invitrogen Cat. #A-11001), protected from light. Plate was then washed 3X with 1X PBS protected from light, counterstained with 300nM DAPI staining solution (Thermo Fisher Cat. #D1306), and then imaged at 10X objective with an Echo Revolve fluorescence microscope.

### 4.16. Quantification and Statistical Analysis

All software used was described in the main text or the appropriate methods section. Statistical tests, as well as statistical comparisons between groups, for each figure were denoted in the corresponding figure legend. P-values for each statistical test were noted in each figure panel, and (adjusted) P-values of 0.05 or lower were considered significant. Analyses were performed in Python and R, using a combination of geomux, cellranger, ScaleAtac, Seurat, archR, CellProfiler, tidyverse, ggplot2, ggpubr, ggrepel, dplyr, tidyr, gridExtra, cowplot, patchwork, stringr, igraph, ggforce, ComplexHeatmap, rstatix, cvequality, EnhancedVolcano.

#### 4.16.1. In silico analysis of chromatin organizer candidates

In our analysis, we rationalized that intra-tumoral heterogeneity is proportional to inter-tumoral heterogeneity due to the central limit theorem. Take the gene *g* and let the number of RNAs of *g* in every cell be *x*; *x* will have a per-cell distribution of *μ*_*x*_ and *σ*_*x*_. Assuming each bulk sample has *N* cancer cells, a bulk sample consists of individual samples (cells) that follow the central limit theorem (CLT). CLT states that:

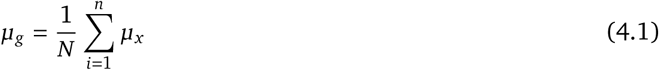

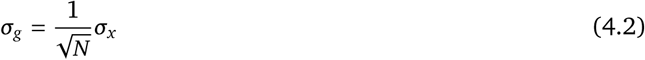

where *μ*_*g*_ is the mean and *σ*_*g*_ is the standard deviation for gene *g* across a population of cells.

Comparing two groups of patient bulk RNA-seq samples may partially reflect differences in underlying cellular heterogeneity between two groups of cells because each group is simply the pooling of multiple patients’ cells. If a factor (gene) such as a chromatin organizer increases *σ*_*x*_ (intracellular heterogeneity), it will also increase the intercellular heterogeneity of *g* across pooled cells (*σ*_*g*_), since *σ*_*g*_ is a function of *σ*_*x*_.

While the CLT predicts attenuation of variance during bulk averaging, factors that increase cellular variance (*σ*_*x*_) should still proportionally increase inter-tumor variance (*σ*_*g*_). Importantly, we acknowledge that inter-tumor variability may also arise from compositional or subtype differences. Therefore, our nomination strategy is intended as a heuristic enrichment approach, subsequently validated using single-cell measurements.

To generate an *in silico* list of chromatin organizer (CO) candidates that could affect transcriptional heterogeneity, we intersected TCGA RNA-seq data with a list of COs collated from Gene Ontology, term *eukaryota mammalia homo sapiens*. Under our *in silico* nomination framework, a chromatin organizer (CO) associated with increased transcriptomic heterogeneity would be expected to stratify tumors such that patients with high CO expression exhibit greater inter-tumor transcriptomic variability than patients with low CO expression.

To implement this, we created a list of patient groupings corresponding to the top and bottom quintile expressers within the set of patients. To reduce the likelihood that our results were biased by a particular set of patients, we also used patient groupings corresponding to the top and bottom quartile expressers. For each patient in the top and bottom grouping, we then calculated genome-wide coefficient of variation (CV; mean-normalized standard deviation) for each available gene, as well as the resulting CV ratio between patients in each grouping. Specifically, we calculated CV across all patients in a particular group, and then calculated the CV ratio for all genes in this manner.

We then generated a p-value for each gene by applying CVEquality for each gene in the given RNA-seq data and converted this to a q-value with a significance cut-off of *α* < 0.05. At this stage, we also recorded the direction of CV change when comparing the groupings, i.e., whether increased expression of a given CO led to increased CV across the given patient set (concordant) or the reverse (discordant). For each CO grouping, we then recorded the number of significant genes, defined as the number of genes flagged as significant via CVEquality testing.

To model the noise inherent to this analysis and derive empirical estimates for the number of genes that would arise by chance via this grouping approach, we generated an empirical null distribution in parallel by randomly sampling tumors and assigning them into groupings of equal size to the above analysis (i.e., quartile and quintile). We used these random groupings of patients (number of groupings used, *n*_groupings_ = 50) to ask how many genes would appear significant if one were to use any given gene to create patient groupings.

Re-applying the computational pipeline described above allowed us to compute estimates for *μ*_random_ (mean number of significant genes arising by chance) and *σ*_random_ (standard deviation of the number of significant genes arising by chance). For each of the CO groupings above, we used these two scalars to compute empirical z-scores for each CO:

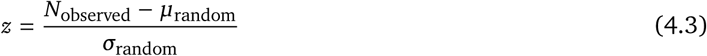

We then applied a z-score cutoff (*z* > 2.5), focusing only on COs above this threshold. For interpretability, we chose to study COs whose higher expression was associated with increased transcriptomic CV. Overlapping nominated COs derived from both quartile and quintile groupings yielded 41 COs that fulfilled our z-score cutoff.

#### 4.16.2. CNA analysis

CNA scores were obtained for TCGA patients to compare differences in CNA profiles of patients in q25 and q75 patient cohorts vs those observed in RNA-seq profiles. Ratios of mean CNA change across patients for each gene was calculated between the two cohorts alongside ratios for mean expression and plotted to observe any correlation present using Pearson’s correlation coefficient.

#### 4.16.3. Tumor purity scores

Tumor impurity was proxied using ESTIMATE scores with higher scores corresponding to higher cell impurities such as immune cells in patient samples. ESTIMATE scores for patients in the quartile heterogeneity analysis were obtained from TCGA to compare differences in tumor purity between q25 and q75 patient cohorts.

#### 4.16.4. PCA heterogeneity analysis

For patient data and cell painting data, we utilized spread of samples (patients or cells) in PCA space as a metric for heterogeneity. For patient data, the spread accounted for transcriptional heterogeneity between TCGA RNA-seq profiles of lower (q25) and higher (q75) RNF8 (or MIS18A in a parallel analysis) expressing patients. Patient’s RNA-seq profiles were put into PCA space and distances of patients in a given quartile (or decile) group from their group’s centroid were calculated and plotted.

For cell morphology, an array of morphology features that CellProfiler measures differences in were assessed. The cell body and ER/Golgi of cells in the non-targeting Ci, RNF8-Ci, MIS18A-Ci, non-targeting Ca, RNF8-Ca, MIS18A-Ca cell painting slides were examined for differences in their shape and volume (see properties in supplemental sheet for features). Cell body and ER/Golgi quantitative property measurements of individual cells were put into PCA space separately, and distances of cells in PCA space from centroid within a sample of cells was calculated for both. Heterogeneity of each sample was measured as the median distance of cells in a sample from the centroid of those cells (centroid distances of cells in the cell body and ER/Golgi PCAs were combined and plotted as boxplots).

#### 4.16.5. Single cell CRISPRi/a RNA-seq

Cell Ranger 7.1.0 software (10x Genomics) was used in the alignment of scRNA-seq reads to the transcriptome, alignment of sgRNA reads to the library, collapsing reads to UMI counts, and cell calling. The 10x Genomics GRCh38 version 2020-A genome build was used as the reference transcriptome. Geomux was used to assign sgRNAs to cells. A hypergeometric distribution was used to calculate the chance of the observed UMI count of sgRNA in cells; default parameters (except min_UMI = 2) and thresholds were used with cells assigned a guide confidently being kept for further analysis.

UMI count data was then normalized using the relative counts method in Seurat and genes with median relative counts of >= 0.1 were filtered for analysis. An equal number of cells were sampled for each guide (n = 90 CRISPRi and n = 70 CRISPRa due to different numbers of cells recovered) to make sure cell count difference did not contribute to gene CV differences across guides. CVs and means were then calculated per CRISPRi/a guide for each gene. Subsequently, in parallel, cell identity (except non-targeting) control was randomized across all cells to create a ‘background’ distribution of expected heterogeneity (shown in **Figure S2A-B**). Individual gene relative counts between guides were plotted as examples to illustrate differences in heterogeneity.

Additionally, to look for differences in cell cycle between guides, cell cycle scores were assigned to cells by the CellCycleScoring function in Seurat and plotted.

#### 4.16.6. Qian et al. single cell dataset

Patients (n = 13) in single cell RNA-seq dataset were grouped by expression levels of RNF8 and MIS18A based on expression quartiles (q25 and q75). Rounding up to whole numbers, 4 patients were selected each for the q25 and q75 cohorts. Data was normalized using the relative counts method in Seurat. Genes with median relative counts of >= 0.1 were filtered for analysis and CV was calculated for these genes and plotted as box plot.

#### 4.16.7. Single cell ATAC-seq

Scale’s Nextflow pipeline was used to align and quantify 10x reads as recovered fragments and assign them to cells as well as assign cells to sgRNAs. The quantified fragments were then analyzed with the archR package’s standard pipeline. The gene score matrix, which is automatically created upon inputting fragment files into archR, was used to assess gene accessibility values. Gene scores are calculated as a function of the placement of a fragment either within the gene or a lower weighted value depending on the distance >5kb from the transcription start or termination site. This is a good proxy for quantifying gene accessibility differences between guides.

An equal number of cells were sampled for each guide (n = 8000) to make sure cell count difference did not contribute to gene CV differences across guides. CVs and means were then calculated per CRISPRi/a guide for each gene from the gene accessibility values. Example gene tracks were constructed using archR. CVs of individual 500 bp tiles (columns in gene track) highlighted in archR gene tracks were calculated across cells (rows in gene track) based on the presence (1) or absence (0) of fragments. A chi-square test was performed using the number of cells with presence or absence of fragment in highlighted tile in cells from guide of interest vs non-targeting control sgRNA cells.

#### 4.16.8. CellProfiler cell morphology for cell painting

Images from high content microscopy at 10x zoom were analyzed using CellProfiler software. The ER/golgi and cell body were examined for properties related to shape and intensity. Quantified cell properties were then analyzed in R using the PCA heterogeneity analysis above.

## 5. Materials availability

All unique/stable reagents generated in this study are available from the lead contact with a completed materials transfer agreement.

## 6. Data and Code availability

The code used in this paper is available at https://github.com/ssobt/nbs_txnheterogeneity. The txnheterogenity package for quantifying heterogeneity between samples in bulk RNA-seq, scRNA-seq, and scATAC-seq data is available at https://github.com/ssobt/txnheterogeneity. Any additional information required to reanalyze the data reported in this paper is available from the lead contact upon request.

## 7. Acknowledgments

We thank Chiara Ricci-Tam, Brian Plosky, Hamed S Najafabadi, Rohit Bose and Luke Gilbert for their invaluable input regarding graphical design, figure layout, and manuscript writing. Sequencing was performed at the UCSF Center for Advanced Technologies (CAT). H.G. is a Core Investigator at Arc Institute, and this research was partially funded by Arc Institute.

## 8. Author contributions

H.G. conceived the study. B.J.W., H.Y., K.G., S.Z. and A.B. designed and performed experiments for the study. S.S. and B.J.W. performed computational discovery of putative chromatin organizers, analyzed experiment data and designed figures. B.J.W. and S.S. wrote the manuscript.

## 9. Competing interests

The authors declare no competing interests.

## Supplementary Material

### A. Supplementary figures

**Supplemental Figure 1, related to Figure 1.**
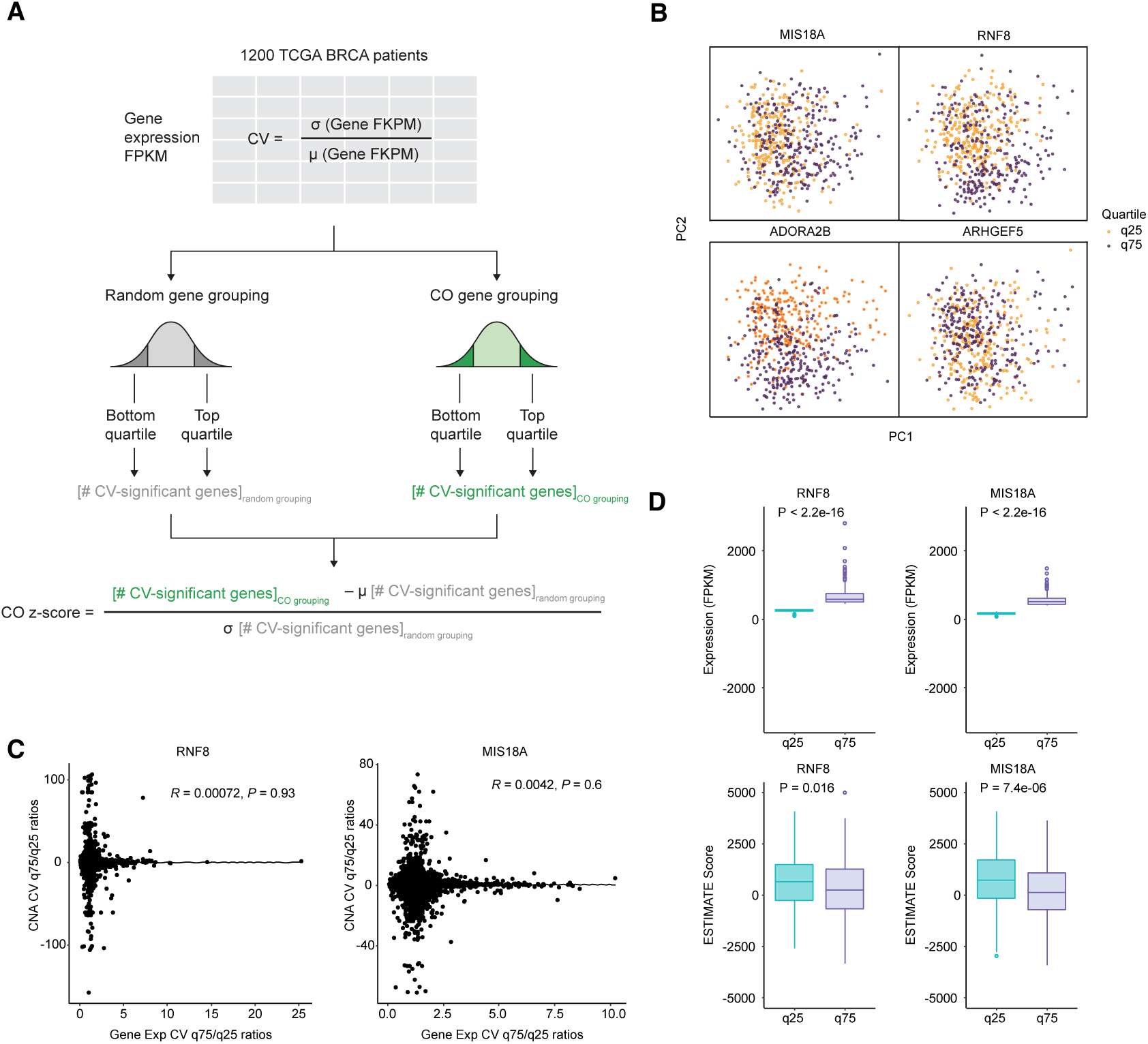
Computational discovery of chromatin organizers correlated with global transcriptomic variation. **(A)** Schematic of the TCGA RNA-seq CV analysis. For each CO, patients were stratified into top and bottom expression quartiles, and genome-wide transcript CV was compared between groups. An empirical null was derived from random gene groupings; COs with normalized *z*-score > 2.5 were selected for further analysis. **(B)** First two principal components for bottom- and top-quartile patients grouped by the indicated genes. **(C)** Comparison of CNA CV ratios and expression CV ratios for genes analyzed in Figure 1B. The two measures are not correlated, indicating that the observed transcriptomic variability is not explained by corresponding copy-number changes. *P*-values by two-sided Student’s *t*-test. **(D)** Top: RNF8 and MIS18A expression levels across bottom- and top-quartile patients. Bottom: ESTIMATE tumor purity scores across the same groups. High purity was observed in both groupings, ruling out confounding by shifts in cellular composition. *P*-values by two-sided Wilcoxon signed-rank test.

**Supplemental Figure 2, related to Figure 3.**
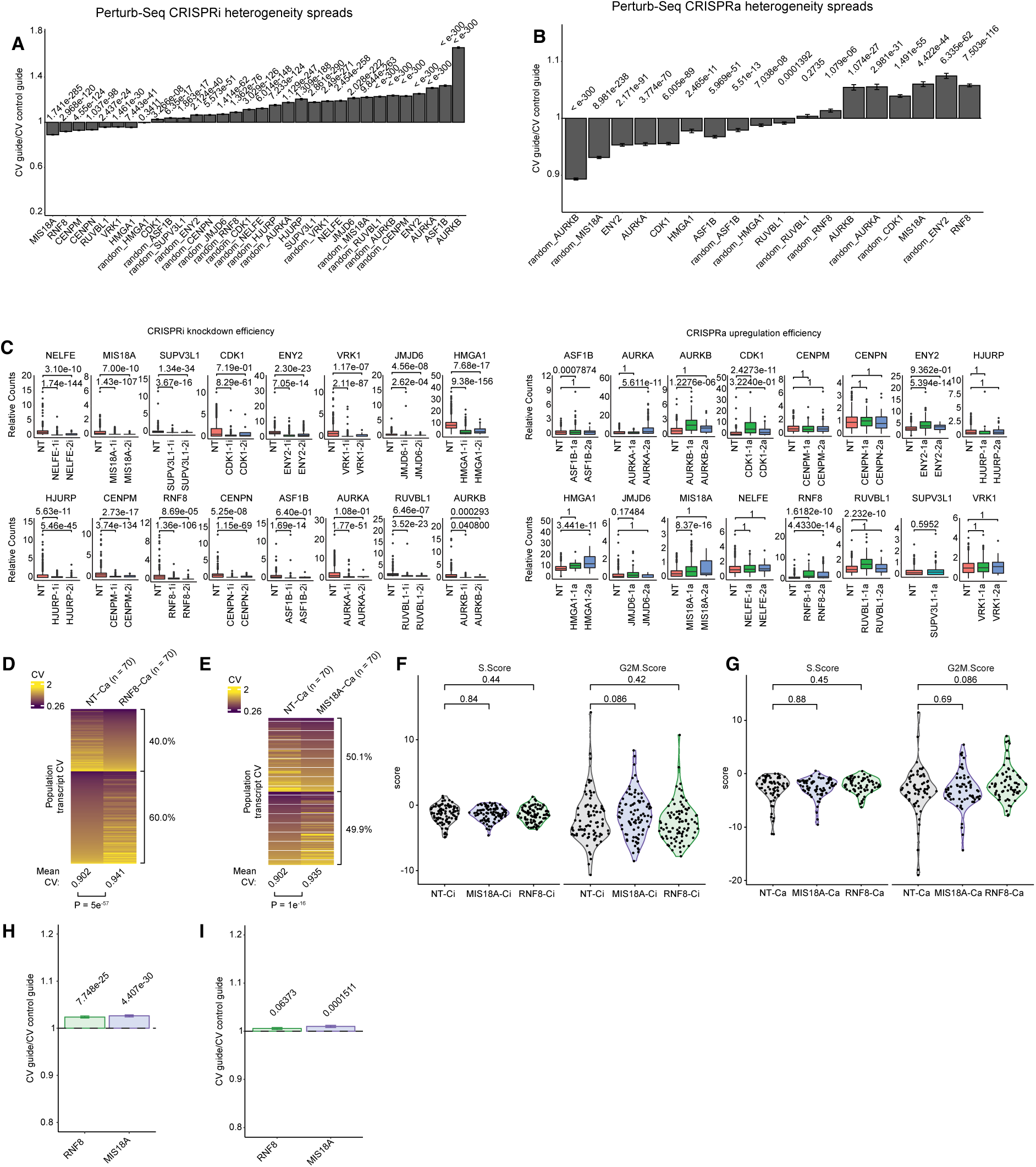
Modulating expression of nominated COs tunes cell-to-cell transcriptomic variability. **(A, B)** Mean and standard error of mean (SEM) barplots for genome-wide CV change upon CRISPRi **(A)** or CRISPRa **(B)** perturbation of all 16 *in vivo*-validated COs. A value of 1 indicates no change relative to non-targeting control. Size-matched random cell groupings serve as empirical null. *P*-values by one-sample two-tailed Student’s *t*-test. **(C)** Per-sgRNA CV change for each guide in the CRISPRi and CRISPRa libraries. Middle and right bars show the two individual sgRNAs per gene. *P*-values by two-tailed Student’s *t*-test, adjusted with the Bonferroni procedure. **(D, E)** Per-gene transcript CV in CRISPRa cells versus non-targeting controls for RNF8 **(D)** and MIS18A **(E)**. Each horizontal line represents a gene; *P*-values by two-sided paired Student’s *t*-test. Percentages indicate proportion of genes that decrease (top) or increase (bottom) in CV relative to non-targeting control. **(F, G)** Cell-cycle phase distribution of cells with the indicated sgRNA perturbation in CRISPRi **(F)** and CRISPRa **(G)** lines. Neither RNF8 nor MIS18A perturbation significantly altered cell-cycle composition. *P*-values by two-sided Wilcoxon signed-rank test. **(H, I)** Mean of the genome-wide gene expression means for the indicated sgRNA perturbations in CRISPRi **(H)** and CRISPRa **(I)** with standard error of mean (SEM) shown. *P*-values by one-sample two-tailed Student’s *t*-test, adjusted with the Benjamini–Hochberg procedure.

**Supplemental Figure 3, related to Figure 4.**
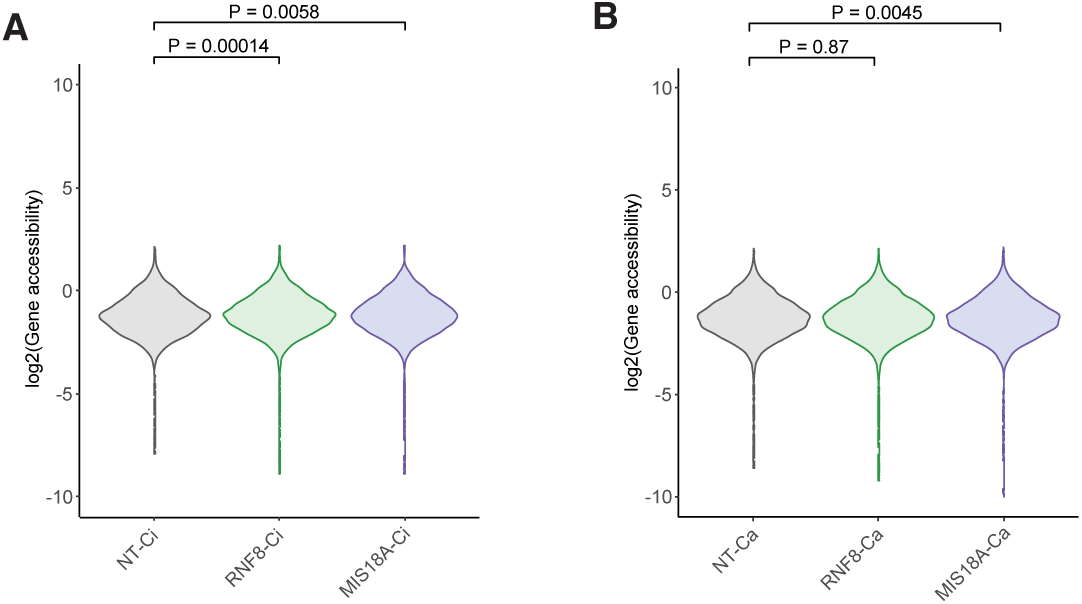
Global mean chromatin accessibility is unchanged upon RNF8 or MIS18A perturbation. **(A, B)** Genome-wide mean chromatin accessibility (pseudo-bulked across cells) for CRISPRi **(A)** and CRISPRa **(B)** perturbations. Accessibility scores were averaged across all assayed gene regions. Neither RNF8 nor MIS18A perturbation shifted mean accessibility, indicating that the observed changes in accessibility variance are not driven by a global shift in chromatin state. *P*-values by two-tailed Student’s *t*-test.

**Supplemental Figure 4, related to Figure 4.**
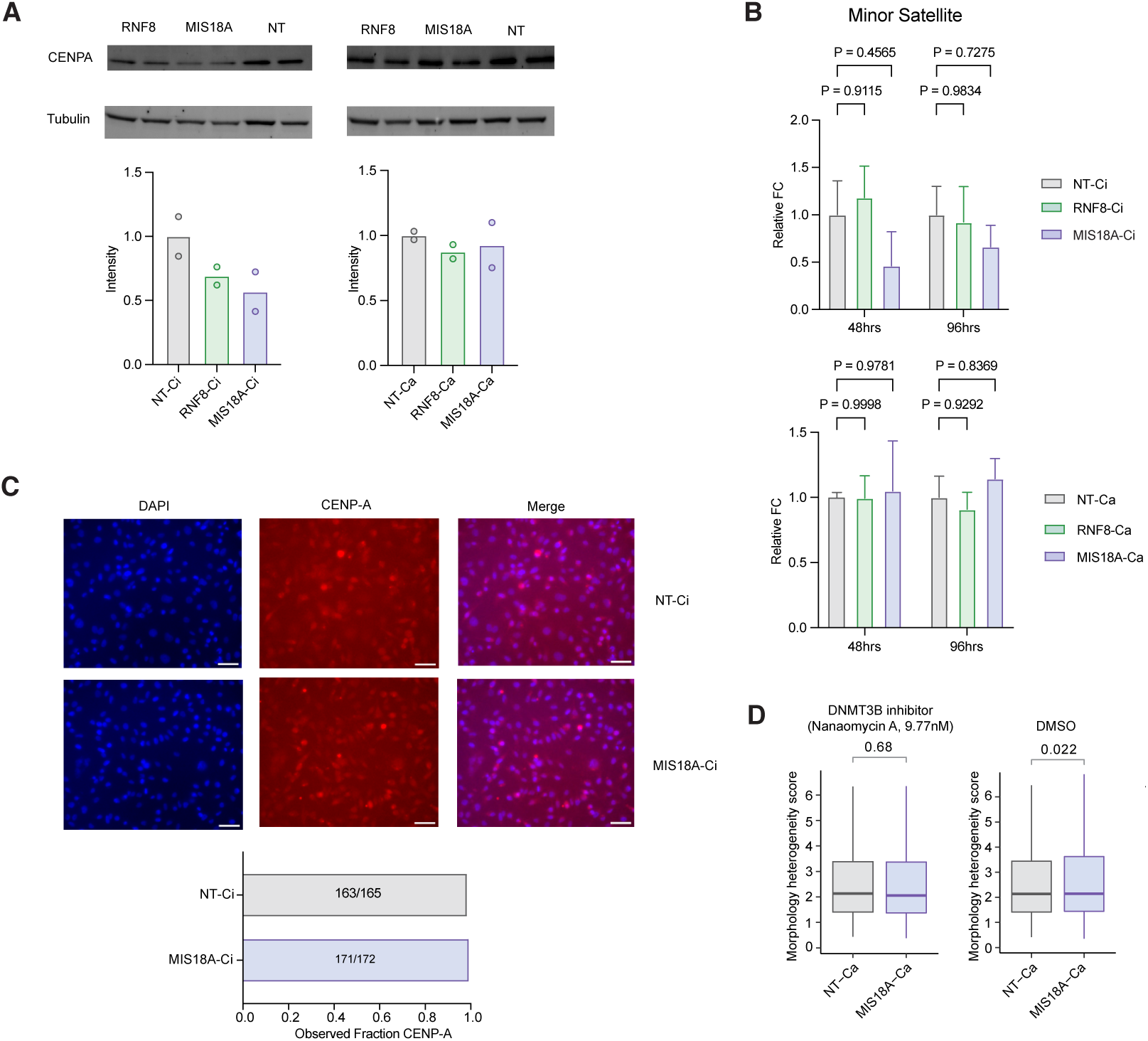
MIS18A perturbation does not disrupt centromeric function. **(A)** Immunoblot of CENP-A in MDA-MB-231 CRISPRi/a lines. Top: CENP-A normalized to tubulin loading control. Bottom: quantification of tubulin-normalized CENP-A signal relative to non-targeting control. **(B)** RT-qPCR of minor satellite ncRNAs in MDA-MB-231 CRISPRi/a lines at 48 and 96 hours post-seeding (*n* = 4). Values are GAPDH-normalized. No significant accumulation of minor satellite ncRNAs was observed upon MIS18A knockdown, indicating that centromeric histone modifications remain intact. **(C)** CENP-A immunofluorescence in MDA-MB-231 CRISPRi/a lines. Top: representative images of CENP-A and DAPI signal. Bottom: quantification of CENP-A as a fraction of DAPI-identified clusters. MIS18A knockdown did not reduce CENP-A signal. Scale bar, 50 *μ*m. **(D)** Cell-painting morphology heterogeneity scores for MDA-MB-231 CRISPRa lines with the indicated sgRNAs, with or without the DNMT3B inhibitor nanaomycin. Nanaomycin rescued MIS18A-overexpression-dependent morphological changes, indicating that MIS18A affects cellular phenotype through DNMT3B-mediated methylation. *P*-values by one-sided Wilcoxon signed-rank test.

**Supplemental Figure 5, related to Figure 5.**
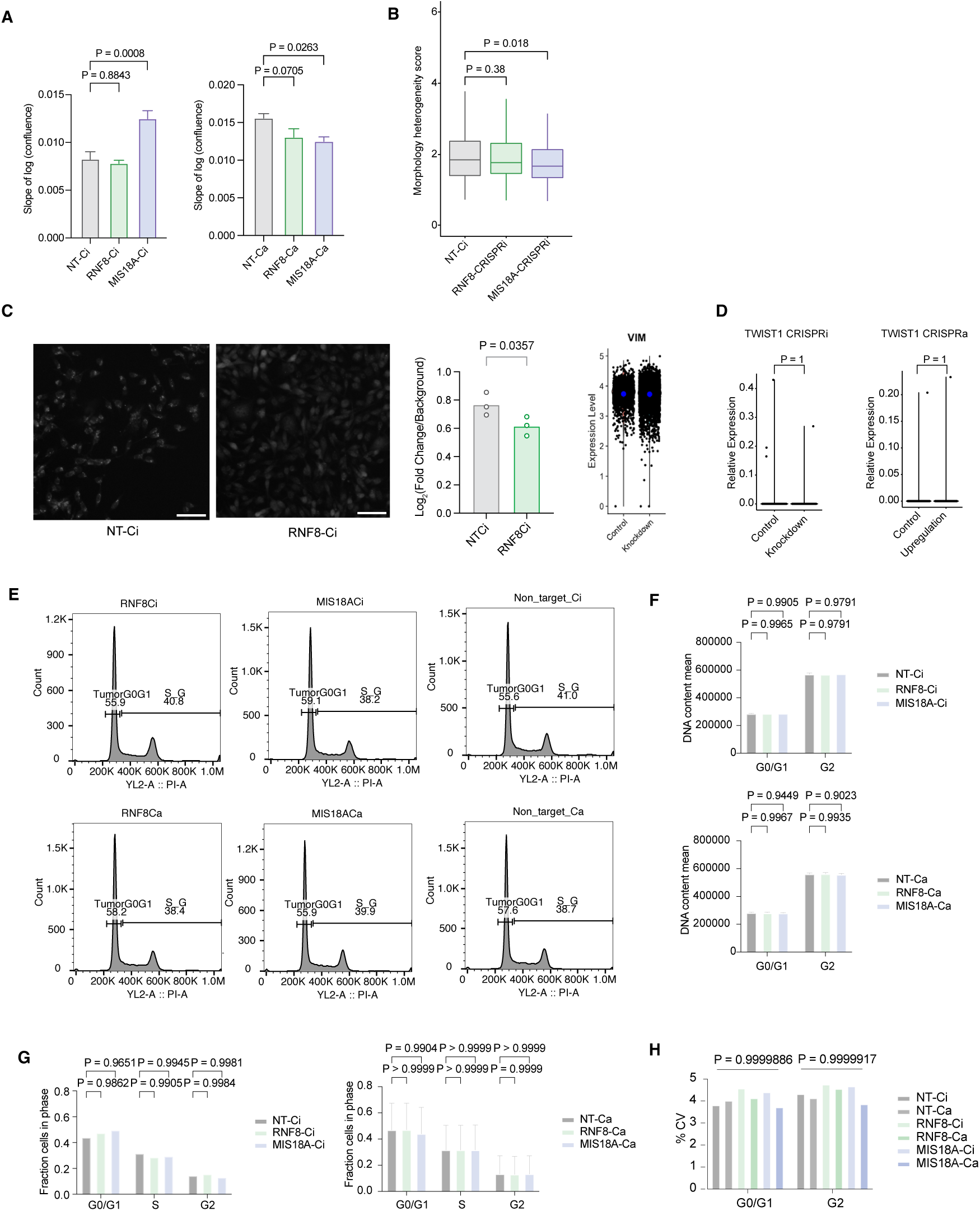
Proliferative, morphological, and genomic characterization of MDA-MB-231 CRISPRi/a lines. **(A)** Cell proliferation rates measured by real-time microscopy. *P*-values by one-way ANOVA (*n* = 8 per condition). **(B)** Cell-painting morphology heterogeneity scores for CRISPRi lines. Scores integrate nuclear, mitochondrial, cell-body, and ER/Golgi features (see Methods). Lower scores (reduced morphological heterogeneity) were observed upon knockdown of either RNF8 or MIS18A. *P*-values by one-sided Wilcoxon signed-rank test. **(C)** Vimentin immunofluorescence in RNF8 CRISPRi lines. Left: representative images showing reduced vimentin signal upon RNF8 knockdown. Scale bar, 68 *μ*m. *P*-value by one-sided *t*-test. Right: vimentin transcript levels from scRNA-seq data in non-targeting versus RNF8 CRISPRi cells; no significant transcriptional change was observed. **(D)** TWIST1 mRNA expression in non-targeting versus CRISPRi (left) or CRISPRa (right) lines. EMT-associated morphological changes were not accompanied by shifts in TWIST1 transcript levels. **(E)** Flow cytometry histograms of propidium iodide-stained CRISPRi/a lines. G0/G1 peaks were called using spiked-in human PBMCs as internal control. **(F)** Mean DNA content for G0/G1 and G2 peaks in each CRISPRi/a line. *P*-values by one-way ANOVA comparing each line to its respective control (*n* = 6). **(G)** Cell-cycle phase fractions for each CRISPRi/a line. *P*-values by one-way ANOVA (*n* = 6). **(H)** DNA content CV for each cell-cycle phase peak. *P*-values by one-way ANOVA (*n* = 6).

**Supplemental Figure 6, related to Figure 6.**
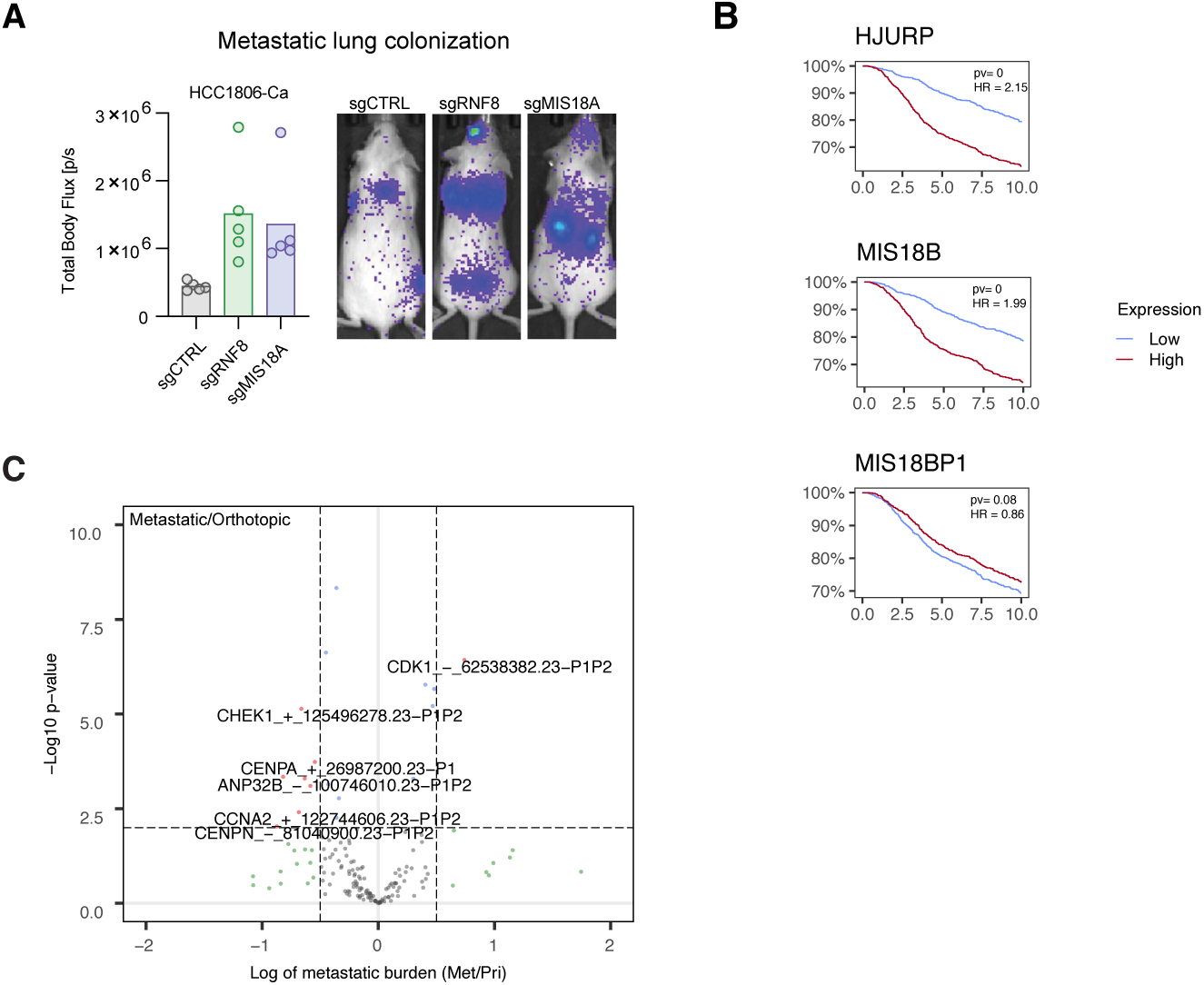
*In vivo* validation and clinical context for RNF8 and MIS18A. **(A)** HCC1806 *in vivo* orthotopic growth assay. Overexpression of either RNF8 or MIS18A increased tumor burden relative to non-targeting controls (*n* = 5 per sgRNA condition). **(B)** Kaplan–Meier survival curves for select CENP-A deposition pathway components, stratified by bottom-quartile (red) versus top-quartile (blue) expression in METABRIC. Higher expression of HJURP and MIS18B was associated with shorter survival; MIS18BP1 showed no significant association. These analyses provide clinical context but do not establish that each complex member modulates ITH. *P*-values by log-rank test. **(C)** Volcano plot of *in vivo* CRISPRi screen results showing metastatic burden normalized to primary tumor burden (tail-vein/orthotopic ratio).

## Notes

### Competing Interest Statement

The authors have declared no competing interest.

